# Neighbor-specific gene expression revealed from physically interacting cells during mouse embryonic development

**DOI:** 10.1101/2021.12.02.470916

**Authors:** Junil Kim, Michaela Mrugala Rothová, Esha Madan, Siyeon Rhee, Guangzheng Weng, António M. Palma, Linbu Liao, Eyal David, Ido Amit, Morteza Chalabi Hajkarim, Andrés Gutiérrez-García, Paul B. Fisher, Joshua M. Brickman, Rajan Gogna, Kyoung Jae Won

## Abstract

Development of multicellular organisms is orchestrated by persistent cell-cell communication between neighboring partners. Direct interaction between different cell types can induce molecular signals that dictate lineage specification and cell fate decisions. Current single cell RNAseq (scRNAseq) technology cannot adequately analyze cell-cell contact-dependent gene expression, mainly due to the loss of spatial information. To overcome this obstacle and resolve cell-cell contact-specific gene expression during embryogenesis, we performed RNA sequencing of physically interacting cells (PIC-seq) and assessed them alongside similar single cell transcriptomes derived from developing mouse embryos between embryonic day (E) 7.5 and E9.5. Analysis of the PIC-seq data identified novel gene expression signatures that were dependent on the presence of specific neighboring cell types. Our computational predictions, validated experimentally, demonstrated that neural progenitor (NP) cells overexpress *Lhx5* and *Nkx2-1* genes, when exclusively interacting with the definitive endoderm (DE) cell. Moreover, there was a reciprocal impact on the transcriptome of the DE cells, as they tend to overexpress *Rax* and *Gsc* genes when in contact with the NP cells. Using individual cell transcriptome data, we formulated a means of computationally predicting the impact of one cell type on the transcriptome of its neighboring cell types. We have further developed a distinctive spatial-tSNE to display the pseudo-spatial distribution of cells in a 2-dimensional space. In summary, we describe an innovative approach to study contact-specific gene regulation during embryogenesis with potential broader implication in other physiologically relevant processes.

**Significance:** Physical contact between neighboring cells is known to induce transcriptional changes in the interacting partners. Accurate measurement of these cell-cell contact based influences on the transcriptome is a very difficult experimental task. However, determining such transcriptional changes will highly enhance our understanding for the developmental processes. Current scRNAseq technology isolates the tissue into individual cells, making it hard to determine the potential transcriptomic changes due to its interacting partners. Here, we combined PIC-seq and computational algorithms to identify cell-type contact dependent transcriptional profiles focusing on endoderm development. We have computationally identified and experimentally validated specific gene expression patterns depending upon the presence of specific neighboring cell types. Our study suggests a new way to study cell-cell interactions for embryogenesis.

## Introduction

Cell-cell contact is important for cell-fate specification during development (1–9). Cell-cell communication including direct cell-cell contact choreographs embryonic patterning, cell-type specification and organ formation (6). Removing specific tissue or placing ectopic explants into various embryonic regions can dynamically change the fate of cells adjacent to the manipulated area (3). For instance, manipulation with the crucial signaling center, anterior visceral endoderm (AVE), causes defects in specification of forebrain identity later in development (10). It is well-recognized that ectopic grafts of signaling centers such as the node or Spemann organizer can induce a secondary neural axis (11, 12). Even with the importance of various cell type interactions for development, it is still challenging to study gene expression changes in association with neighboring tissue. While the grafting experiments have significantly enhanced our understanding of the inductive ability of various neighboring cell types (11), these experiments are technically demanding and/or are not always feasible.

Single cell transcriptional profiling has been successfully applied to identify cell types and their developmental trajectory during mouse organogenesis (13–15). However, the loss of spatial information in scRNAseq makes it difficult to trace the genes induced and regulated by interaction of different cell types. Co-expression of ligand-receptors pairs from scRNAseq data has been used to predict interacting cell types (16, 17). However, there are diverse ways of cell communication besides ligand-receptor, such as direct cell communication through gap junctions (18). It is still difficult to accurately detect neighboring tissue-specific gene expression changes.

Recently, RNA sequencing of multiple-interacting cells has been used to measure the transcriptome of physically interacting cells (PICs) without relying on ligand-receptor co-expression (19–23) ProximID identified the interacting cell types in the bone marrow and the small intestine in mice using the transcriptome of two or more PICs by applying mild dissociation of cells (19). PIC-seq was employed to interrogate interactions of immune and epithelial cells in neonatal murine lungs (22) and to understand transcriptomic changes of liver endothelial cells (LECs) across liver zones (20). Cell-cell interactions by multiple sequencing (CIM-seq) was applied to identify interacting cell types in gut epithelium, lung, and spleen (23).

While previous approaches using multiple interacting cells have mainly focused on investigating interacting cell types, we hypothesize that interacting multi cells will retain gene expression derived from the physical interaction defined by these disparate cell types. To identify cell-type contact associated transcriptional profiles focusing on endoderm development, we dissected the developing mouse embryos (at E7.5, E8.5 and E9.5), sorted FOXA2^Venus^ expressing cells and used this material for PIC-seq. Using PIC-seq, we captured both homotypic and heterotypic cell clumps physically interacting with each other. Interestingly, we identified various sets of genes expressed in the heterotypic PICs but not each individual cell type. Computational analysis identified the cell types expressing unique gene sets specific to their neighbors. For instance, *Lhx5* and *Nkx2-1* were expressed exclusively in the NP cells that physically interact with DE cells and *Gsc* and *Rax* were expressed in the DE cells interacting with NP cells. Some of these neighboring cell type specific genes were associated with the development of specific embryonic regions. Validation using Geo-seq (24) which provides transcriptome from the dissected developing mouse embryos confirmed that we had successfully identified spatially organized sets of cells and neighboring cell specific genes. Additionally, we designed experimental tools to investigate and successfully validate the computational predictions. Notably, we were able to predict neighboring cell types from scRNAseq based on the cell-contact-specific genes. We further emulated the anatomy of the mouse embryos by visualizing cell-cell contact information and neighboring cell specific gene expression in a modified t-distributed stochastic neighbor embedding plot (spatial-tSNE). Our results suggest that the local environment information retained in the transcriptome of individual cells can be used to reconstruct potential spatial gene regulation patterns during development.

## Results

### Mapping of PIC-seq data onto cell types in the mouse embryo

To identify transcriptional profiles influenced by cell-cell contact during development with focus on the endoderm, we took advantage of a mouse line in which a Venus fluorescent protein had been fused to the endoderm regulator FOXA2 to identify FOXA2 expressing endoderm cells from embryos ranging from E7.5 - E9.5 (Fig. 1A). We obtained PIC-seq after mild dissociation of these mouse embryos followed by fluorescence-activated cell sorting (FACS) against FOXA2^Venus^, and cell size was used to filter out potential single cells.

**Figure 1.**
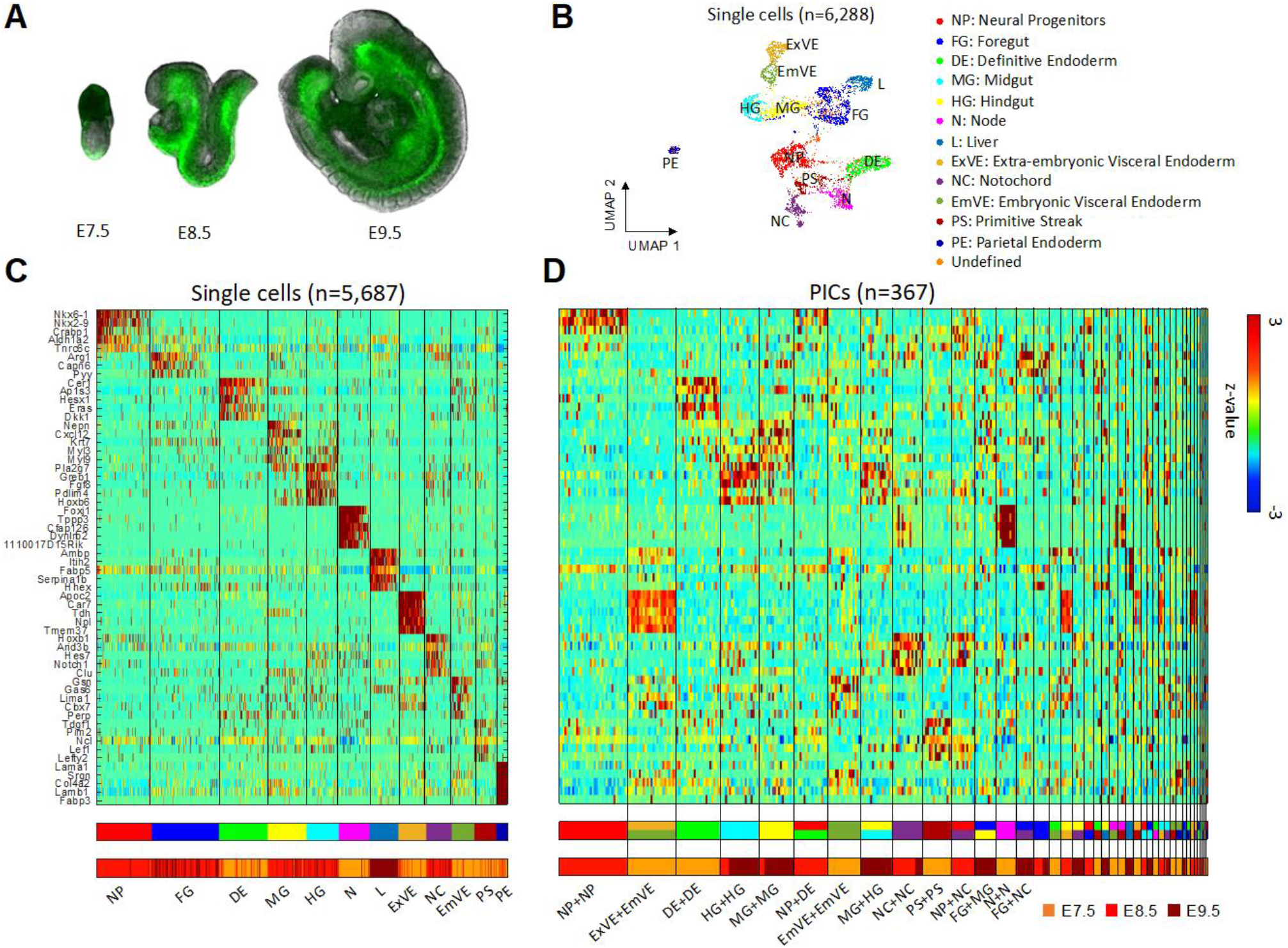
PIC-seq for developing mouse embryos. (**A**) A cartoon of mouse embryos (E7.5, E8.5 and E9.5) with fluorescently tagged FOXA2^Venus^. (**B**) Clustering analysis identifies 12 cell types from scRNAseq data from the developing mouse embryos. (**C**) A heatmap of the marker genes exhibits distinct expression profiles for the 12 identified cell types. The cell types from the clustering analysis and the embryonic days are shown at the bottom (**D**) Gene expression profiles of the identified marker genes for 367 PICs. The predicted cell types for the PICs are shown and their embryonic days are shown at the bottom.

After applying stringent quality control measures, RNA sequencing data from 367 PICs were obtained using massively parallel RNA single-cell sequencing (MARS-seq) technology (25, 26). To analyze the PIC-seq data, we used single cell RNA sequencing (scRNAseq) from MARS-seq (total 6,288 cells) against mouse embryos obtained at matching time points (27). Data is presented to summarize the number of PIC-seq pairs and the number of cells from scRNAseq obtained at each stage as well as their distribution of the number of genes and the read counts (Fig. S1).

From the scRNAseq, we identified 13 clusters: neural progenitors (NP; n=767), foregut (FG; n=927), midgut (MG; n=537), hindgut (HG; n=444), definitive endoderm (DE; n=677), liver (L; n=402), extra-embryonic visceral endoderm (ExVE; n=385), embryonic visceral endoderm (EmVE; n=314), primitive streak (PS; n=302), node (N; n=422), notochord (NC; n=350), parietal endoderm (PE; n=160) and one undefined cluster (Fig 1B, 1C) and (Fig. S2). From the cluster analysis, we identified 545 cell-type specific marker genes (FDR < 0.01 & log ratio > 0.4) (Table S1).

To annotate the cell types of the PICs, we used the cell type marker genes identified from scRNAseq clustering (Fig. 1C). For this purpose, multiclass support vector machines (SVMs) were trained with artificial doublets from two randomly selected single cells for the three time points. SVMs were known to perform well for classifying high-dimensional data even with a small number of samples (28). The trained SVMs classified the 367 PICs into homotypic and heterotypic combinations of cell types (Fig. 1D and Table S2). The error rate of 10-fold cross-validation for the SVMs were 17.27%, 18.76%, and 28.56% in E7.5, E8.5, and E9.5, respectively, suggesting comparable performance with a previous deconvolution approach (22). The designated marker genes were well expressed in the annotated PICs (Fig. 1D). Notably, the frequencies of the heterotypic combinations identified using PICs were significantly high as compared with the combination of erroneous doublets identified using DoubletFinder (29) (Fig. S3). The observed heterotypic combinations such as ExVE+EmVE (at E7.5) and NP+DE (at E8.5) reflect the neighboring tissues in developing mouse embryos (13, 30, 31).

### PIC-seq enables detection of cell-cell contact specific gene expression

To examine cell-cell contact-specific gene expression, we investigated genes highly expressed in the heterotypic PICs but not in the homotypic combinations of individual cell type. For the PIC types with at least 10 cell type pairs (ExVE+EmVE in E7.5, NP+DE in E8.5, NP+NC in E8.5, MG+HG in E9.5, and FG+MG in E9.5), we investigated neighboring cell-cell contact-associated gene expression (Figs. 2 and S4-S7). We identified 167 genes highly expressed in the heterotypic PICs as compared to the expected expression levels for individual cell types obtained from scRNAseq (FDR < 0.01 & log ratio > 0.5) (Table S3). For instance, the heterotypic PICs for NP+DE expressed genes such as *Lhx5, Nkx2-1*, and *Gsc*. These PICs also expressed the marker genes for NP and DE: *Crabp1* and *Col8a1* for NP, and *Trh* and *Slc2a3* for DE (Figs. 2A and 2B). Interestingly, some of the identified neighboring cell type specific genes are known to play crucial roles during development. *Lhx5* is known to promote forebrain development (32). *Nkx2-1* is required for the proper specification of interneuron subtypes (33). *Gsc* is a known marker of anterior endoderm (34), pre-chordal plate (35), and has been implicated in ESC differentiation to definitive endoderm differentiation (36–38).

**Figure 2.**
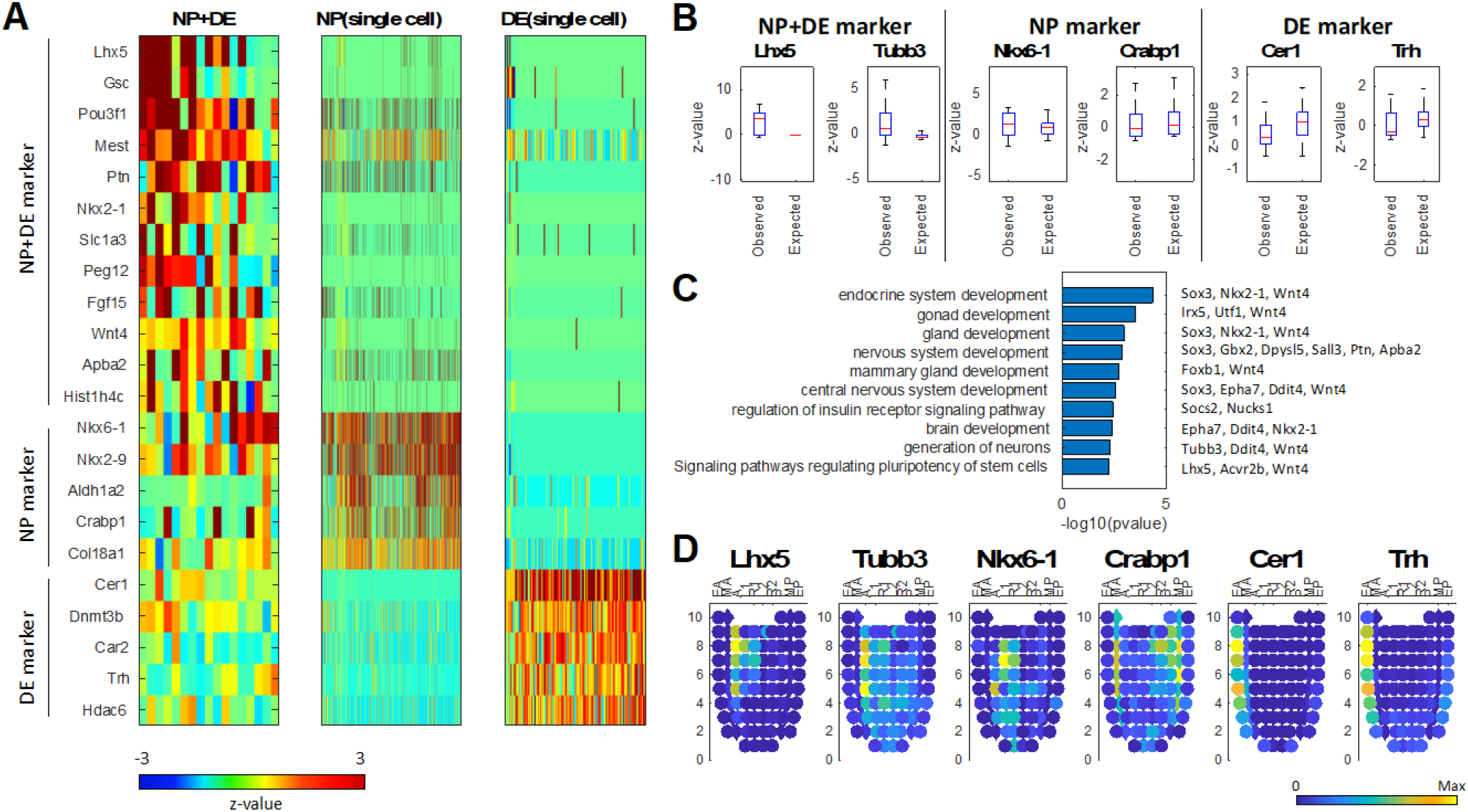
PIC-seq analysis identified genes highly expressed in NP+DE. (**A**) A heatmap of PIC-seq of NP+DE showed contact-specific expression as well as the marker genes for NP and DE. (**B**) Contact-specific expression levels are significantly high for the PIC-seq compared with the expected expression levels obtained from scRNAseq. (**C**) GO and KEGG pathway terms enriched in the contact-specific genes. (**D**) Validation of the contact-specific marker genes using Geo-seq data. EA: Anterior Endoderm; MA: Anterior Mesoderm; A: Anterior; L1: Anterior Left Lateral; R1: Anterior Right Lateral; L2: Posterior Left Lateral; R2: Posterior Right Lateral; P: Posterior; MP: Posterior Mesoderm, EP: Posterior Endoderm.

The 54 genes highly expressed in the heterogeneous PIC of NP+DE at E8.5 were linked to endodermal or neuroectodermal embryo development including gland development (*Sox3, Nkx2-1*, and *Wnt4, p*-value: 1.1E-3) and nervous system development (*Sox3, Gbx2, Dpysl5, Sall3, Ptn*, and *Apba2, p*-value: 1.38E-3) based on the gene ontology (GO) terms and Kyoto Encyclopedia of Genes and Genomes (KEGG) pathways enrichment tests (39) (Fig. 2C). The 39 NP+NC contact-specific genes were mainly associated with mesodermal development, e.g., *Mesp1, Meox1, Lef1* (Fig. S6). Our results suggest that PIC-seq could be used to identify and stratify genes induced in regions where different cell types are in physical contact during embryonic differentiation.

### Contact-specific gene expression is spatially localized at the boundary regions between two cell types

To validate the spatial expression of contact-specific marker genes, we used a publicly available Geo-seq dataset (24) which contains transcriptome from 83 dissected regions in the developing mouse embryos at E7.5 (Figs. S8, S9 and S10). DE marker genes (*Cer1* and *Trh*) were strongly expressed in anterior endoderm (EA), while NP marker genes (*Nkx6-1*, and *Crabp1*) were strongly expressed interior to the endoderm in Geo-seq (Figs. 2D and S8). Intriguingly, the contact-specific genes (*Lhx5* and *Tubb3*) were expressed in-between the regions expressing NP and DE specific genes (Figs. 2D and S9). The averaged profiles for NP+DE genes also showed strongest expression of the contact-specific genes in between NP and DE marked regions in the Geo-seq (Fig. S10A).

In addition, we found that the ExVE+EmVE specific genes were mainly expressed in-between the regions marked by the two cell types (upper part of the corn plot) in the Geo-seq dataset (Fig. S10B). While the nature of the GEO-seq dataset is such that it does not include structures like gut or notochord, we identify co-expression of contact associated genes, e.g. the NP+NC specific genes were highly expressed in the posterior mesoderm (Fig. S10C), which was located between the posterior ectoderm and node region (bottom of the corn plot). Our results indicate that the contact-specific genes obtained from PIC-seq reflect the initial and accurate spatial expression.

To further confirm our observations, we used the public seqFISH data (40) from mouse embryos at E8.5-E8.75 for 351 genes. The cell type annotation (brain/spinal cord, gut tube, and DE) visualized the anatomical structure for NP, DE, FG, MG, and HG (Fig. S11A). The NP markers (*Aldh1a2, Sfrp1*, and *Sox2*) and the DE markers (*Cdh1, Cer1, Dkk1, Dnmt3b, Krt18, Otx2, Sfrp1*, and *Sox17*) were highly expressed in the brain/spinal cord and the DE regions, respectively (Fig. S11A). The zoomed-in view at the boundary regions for NP and DE shows that the contact-specific genes (*Foxb1, Gbx2, Irx5, Lefty1, Pou3f1, Ptn, Sall3* and *Socs2*) were expressed in between those two regions (Figs. S11B and S11C).

### Prediction of the neighboring cell type from scRNAseq tells the contributing cell type for the gene expression changes in PIC-seq

Based on the contact-specific gene expression, we originated an approach to predict neighboring cell types by using the transcriptome of single cells. The neighboring cell type can be predicted by examining specific neighbor specific genes detected from PICs. Therefore, the contact-specific genes in the PIC-seq were used to train a multiclass SVM. The trained SVM predicts the interacting cell types of a cell by forming artificial doublets and voting from the SVM output. Among the 767 NP cells, 85 were predicted to neighbor with DE, 123 interacted with NC and the majority (559 cells) interacted with other NP cells (Fig. 3A). Among the 677 DE cells, 252 were predicted to interact with NP (Fig. 3B). Notably, neighboring cell type prediction further informed us of the cell type expressing contact-specific genes. Among the contact-specific genes in the heterotypic NP+DE PICs, *Lhx5, Pou3f1, Mest, Ptn, Nkx2-1*, and *Slc1a3* were from the NP cells and *Gsc, Sox3*, and *Rax* were from the DE cells (Figs. 3A, 3B, and Table S4). These analyses further predicted the list of neighboring cell specific genes expressed in each cell type. For instance, NP cells expressed *Lhx* and *Nkx2-1* when interacting with DE and *Prtg* and *Gas1* when interacting with NC cells (FDR<0.01) (Fig. 3A).

**Figure 3.**
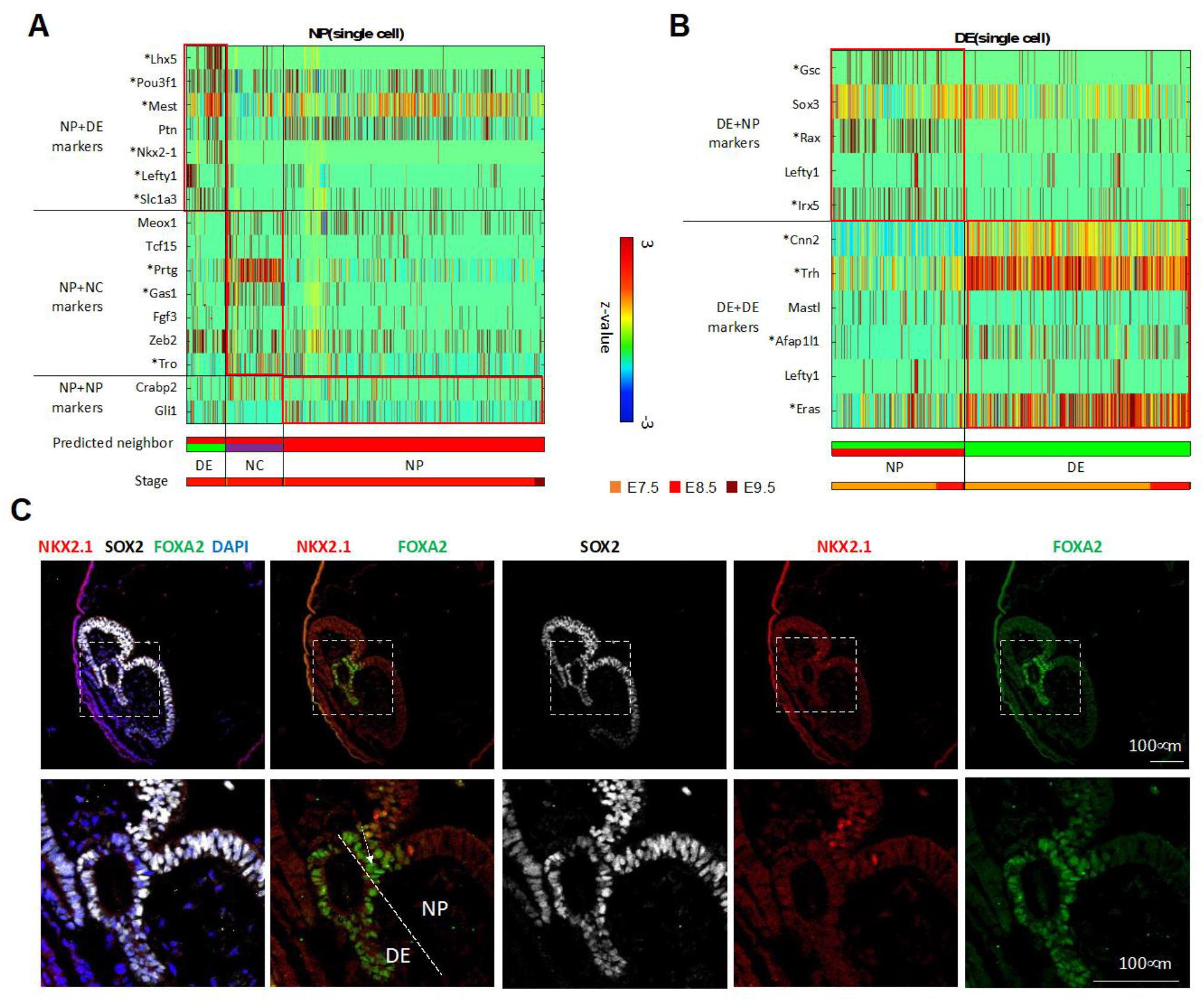
Neighboring cell type prediction from scRNAseq. (**A**) Single NP cells in the original MARS-seq dataset that are predicted to interact with DE or NC. Predictions are based on expression of defined neighbor specific genes from PIC-seq. (**B**) Single DE cells in the original MARS-seq dataset that are predicted to interact with NP. Predictions are based on expression of defined neighbor specific genes from PIC-seq. (**C**) Validation of a contact-specific gene Nkx2-1 using staining of a mouse embryo. Nkx2-1 is expressed in the NP (SOX2+) cells contacting with DE (FOXA2+).

To validate these contact-specific genes, we performed co-staining of FOXA2 (a DE marker), SOX2 (an NP marker), and NKX2-1 (a contact-specific gene) in the mouse embryo at E8.25. FOXA2 was expressed in DE as well as the floor plate that connects to NP. SOX2 was expressed stronger in NP (compared with DE). NKX2-1 was expressed in the part of NP cells when they were proximal to DE, consistent with our prediction (Fig. 3C).

We designed a specific set of experiments to experimentally validate the accuracy of the computational predictions. Focus was on a subset prediction that involves the impact of contact between the NP and DE cells on their respective transcriptome. The experimental strategy we adopted is demonstrated graphically (Fig. 4A). The strategy involves the isolation of NP and DE cells from E8.5 mouse embryos, using pre-defined cell-specific markers, NCAM-1 positive, SSEA-1 positive and SSEA-4 negative for NP cells and CD24 positive, Claudin positive, SSEA-1 negative and SSEA-4 negative for DE cells. The isolated cells were maintained in culture media. A specific set of DE cells were infected with comet-pD2109-CMV lentiviral particles expressing blank green fluorescent protein (GFP) and mixed with NP cells in 1:1 ratio. The GFP positive DE cells and GFP negative NP cells were co-cultured for 48 hrs. At the end of this 48 hrs., the GFP positive DE cells and GFP negative NP cells were sorted using flow-based cell sorter (Fig. 4B). The expression of 4 genes *Lhx5, Nkx2-1, Gsc* and *Rax* was observed in the NP and DE cells maintained as a monoculture and the NP and DE cells that were in contact with each other in a Co-culture experiment (Fig. 4C). The qPCR-based mRNA expression analysis revealed elevated expression levels of *Lhx5* and *Nkx2-1* genes in NP cells exclusively when they were in contact with DE cells. The expression of these genes in the NP cells maintained in monoculture was around 2.8-fold for *Lhx5* and 2.2-fold for *Nkx2-1* lower, when compared to NP cells which came in contact with DE cells. Similarly, the qPCR-based mRNA expression analysis revealed elevated expression levels of *Gsc* and *Rax* genes in DE cells exclusively when they were in contact with NP cells. The expression of these genes in the DE cells maintained in monoculture was around 3.2-fold for *Gsc* and 3.7-fold for *Rax* lower, when compared to DE cells which came in contact with NP cells. This experiment suggests cell-cell contact or local paracrine signaling can induce *de novo* gene expression at different stages of development and our computational methods are able to accurately detect these signals (Fig. 4C).

**Figure 4.**
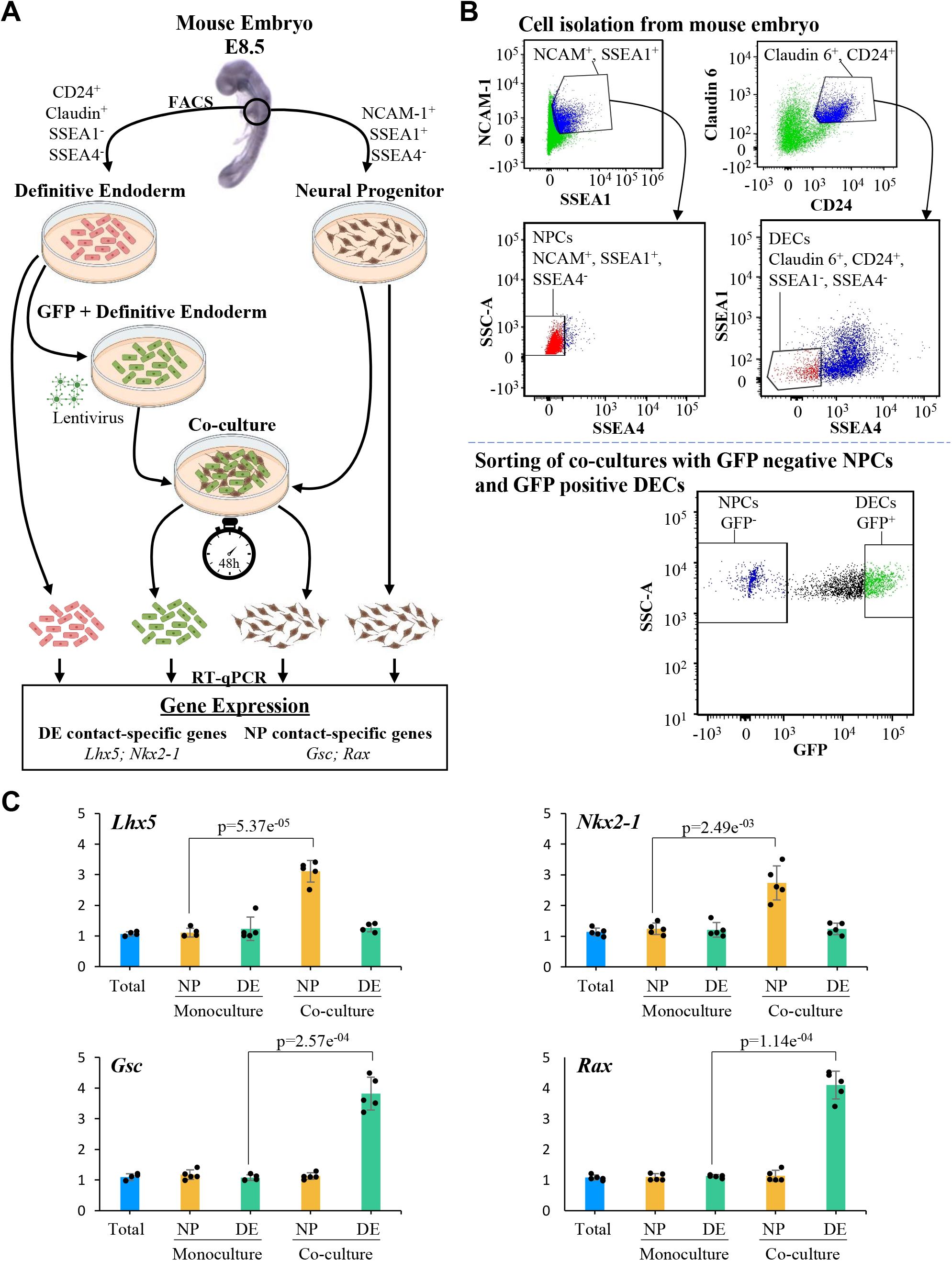
Experimental validation of the neighboring cell type prediction. (**A**) A model depicting the experimental approach used to analyze changes in the expression of contact-specific genes from NP and DE cells. Briefly, we isolated NP and DE cells from E8.5 mouse embryos, tagging DE cells with GFP. Then we performed monoculture of DE and NP cells as well as co-culture of both cell populations for 48 h. After co-culture, DE and NP cells were sorted by GFP expression. Finally, we used monocultured and sorted DE and NP cells to perform RT-qPCR, which allowed us to measure the expression of the DE contact-specific genes *Lhx5* and *Nkx2-1*, and the NP contact-specific genes *Gsc* and *Rax*. (**B**) FACS of NP and DE cells from mouse embryo at E8.5. On the left-top gate, we sorted double positive cells for NCAM-1 and SSEA1 (blue population). From this population we sorted SSEA4 negative cells, which represent NP cells (red population, left-middle gate). On the right-top gate, we sorted double positive cells for Claudin-6 and CD24 (blue population). Then, we sorted SSEA1 and SSEA4 negative cells, to obtain DE cells (red population, right-middle gate). The bottom gate shows the sorting of DE and NP cells after co-culture for 48h. Cells were sorted accordingly to GFP expression since DE cells were tagged with GFP prior to their use in the co-culture experiment. (**C**) The analysis through RT-qPCR of contact-specific genes from NP and DE cells shows that, after co-culture, NP cells upregulate *Lhx5* (*p*=5.37E-5) and *Nkx2-1* (*p*=2.49E-3) while DE cells increase the expression of *Gsc* (*p*=2.57E-4) and *Rax* (*p*=1.14E-4). Expression in these genes in the total embryonic tissue represents the control (lane 1, blue bar in every plot), the relative expression of these genes in the monoculture and co-culture is calculated relative to the expression of these genes in the total embryonic tissue (N=5). We performed a two-tailed Student’s t-test.

We also predicted the neighboring cell type for FG, MG, HG, NC, and EmVE cells using the same strategy (Fig. S12). We further tested if the trained SVM could annotate the neighboring cell types for publicly available scRNAseq data from developing mouse embryos (13, 15). After the annotation, we still found the distinctive groups of DE cells expressing contact-specific genes when contacting NP such as *Rax* and *Gsc* in these independent datasets (Figs. S13 and S14). Our results indicate that scRNAseq retains the information about the neighboring cell type even after cells are isolated. In summary, we observed a diverse repertoire of contact-specific genes depending on their neighboring cell types.

### Visualizing spatio-structure of tissue using spatial-tSNE

Our prediction suggests that the transcriptome of a cell contains information about the neighboring cell type. However, the current visualization algorithms for scRNAseq including UMAP (Fig. 1B) or t-distributed stochastic neighboring embedding (tSNE) cannot accurately represent the neighboring cell types. To visualize neighboring cells and the neighboring cell type specific expression profiles, we revisited the tSNE algorithm which assigns small probabilities when locating cells in the 2D plot for the cells whose transcriptomic distances are large. The spatial-tSNE algorithm we developed can visualize the neighboring cells located near each other by assigning the highest probability for the cell pairs that are classified into neighboring cells. Compared with conventional visualization approaches based on the transcriptomic similarities (Fig. 1B), spatial-tSNE provides information about interacting cell types in the mouse embryos. For instance, the spatial-tSNE showed the physical interactions between EmVE and DE cells (Fig. 5A), while they were distally located in the classical UMAP plot (Fig. 1B). In addition, the spatial-tSNE plot provided spatial layouts of NP+DE, MG+NC and DE+EmVE, which are consistent with the anatomy of the mouse embryos.

**Figure 5.**
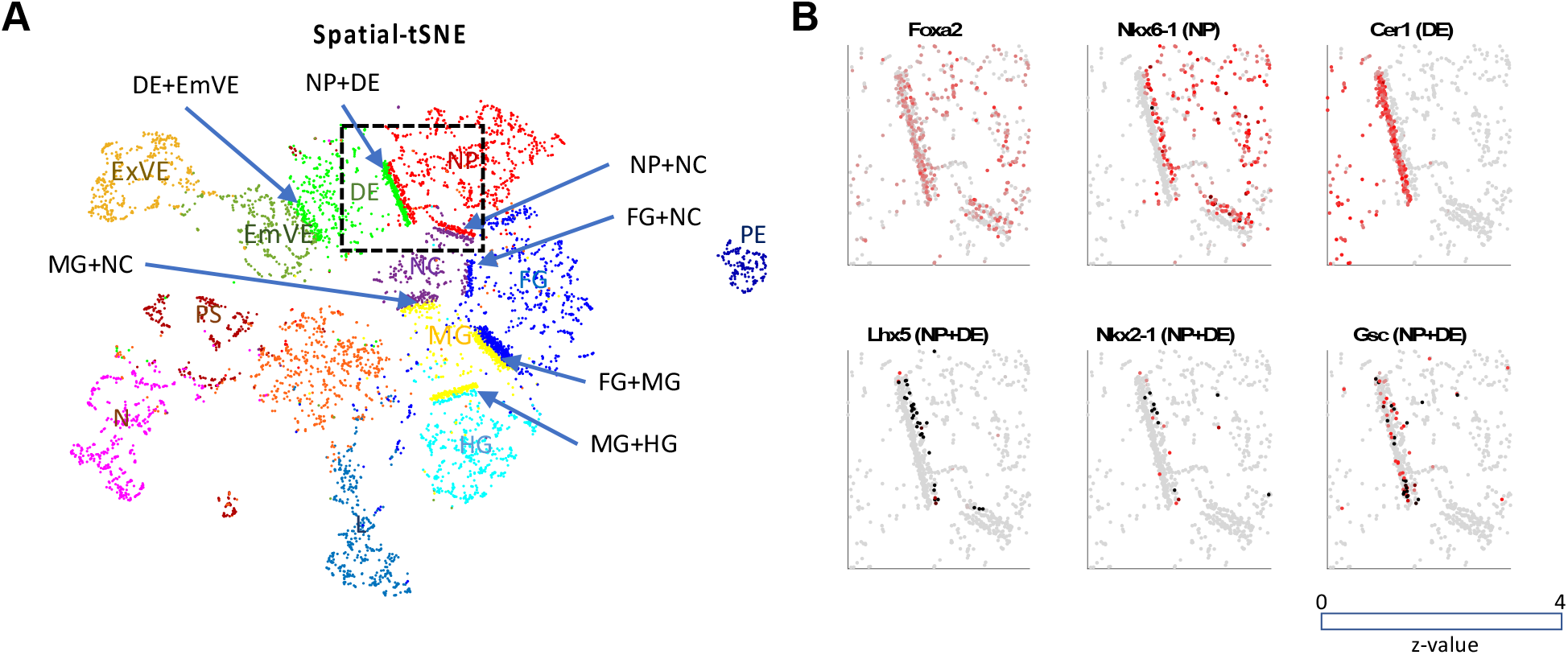
Visualizing spatio-structure of tissue using spatial-tSNE for scRNAseq. (**A**) A spatial-tSNE plot recapitulating the spatial distribution of cells in mouse embryos. (**B**) Expression patterns of NP, DE, and NP+DE markers on the spatial-tSNE plot for the boxed region in A.

The spatial-tSNE plot summarizes the neighboring cell type specific expression patterns in 2D embedded dimensions (Fig. 5A). The average expressions of NP+DE, MH+HG, and NP+NC markers were localized near the border of the corresponding two cell types (Fig. S15). Spatial-tSNE visualizes the neighboring cell-type specific factors. For instance, *Lhx5* is expressed in NP cells close to DE and *Gsc* is expressed more profoundly in DE cells close to NP (Fig. 5B).

## Discussion

Cells are influenced by the neighboring cells in many ways including cell size, stiffness, and mechanical forces (41, 42). During embryogenesis, coordination between adjacent cells is essential to regulate the expression of genes in correct spatial and temporal contexts. To establish these patterns during morphogenesis, these cells influence each other’s gene expression by exploiting various signaling molecules, direct cell-cell contact (43) or reconfiguring mechanical environment (44, 45).

Here we asked if there are unique genes expressed when cells are in contact with each other. To obtain cell-cell contact information, we used PIC-seq and established computational algorithms to identify neighboring cell type specific gene expression. PIC-seq, by retaining cell contact, enabled us to predict neighboring cell type specific gene expression. Our predictions indicated that cells present specific gene expression patterns depending on their neighboring cell type (Fig. 3A). For instance, NP cells expressed *Lhx5* and *Nkx2-1* when neighboring DE and *Gas1* when neighboring with more posterior NC.

To confirm our observation in an unbiased way, we used independent publicly available Geo-seq (24) and seqFISH (40) datasets. The neighbor-specific genes we identified were spatially expressed between the regions marked for each cell type (Fig. 2). Even though Geo-seq (24) is limited to the mouse embryos at E7.5, it was sufficient to highlight the likely spatial location of pairs of interactors for NP+DE, ExVE+EmVE and NP+NC (Fig. S10). The seqFISH data (40) also confirmed that neighbor-specific genes are expressed in the regions between two cell types.

We further questioned if cell-contact or proximity induces specific gene expression. For this, we devised a co-culture system followed by sorting individual cell types and measured gene expression changes (Fig. 4). Our experimental strategy clearly showed that there are genes induced by cell-contact or proximity and PIC-seq provided an approach to obtain this information in an unbiased manner.

Among the predicted neighboring cell type specific genes, we identified several factors with established roles during development. For instance, *Lhx5*, a gene expressed in NP cells when contacting DE, is known to promote forebrain development by regulating the Wnt signaling pathway (32). Knockdown of *Lhx5* resulted in apoptotic hypothalamic development (46). *Nkx2-1*, another gene expressed in NP cells when contacting DE, is also recognized for its role in response to dorsal-ventral patterning in the neural tube and for specifying cortical interneuron subtypes (33). Also, DE cells interacting with NP cells expressed *Gsc* (Figs. 2A and 3B) which has a role in anterior endoderm (34), pre-chordal plate (35), definitive endoderm differentiation (37) and foregut formation (38). Our results indicate that cell contact or proximity has the potential to activate cell type specification during embryogenesis.

Our study indicates that a cell encodes details about its local environment in its transcriptome. For instance, single NP cells identified as interacting with DE retained the expression of *Nkx2-1* even after cell isolation (Figs. 2A and 3A). Based on the local information detailed in the transcriptome of a cell, we were able to predict its neighboring cell type (Figs. 3A and 3B). The trained model was further applied to identifying neighboring cell types for publicly available scRNAseq during embryogenesis (13, 15) (Figs. S13 and S14). This indicates that the contact-specific genes identified from PIC-seq can be used as a reference to re-annotate the neighboring cell types of public scRNAseq.

Gene expression varied between cells as a function of their neighbors. In our study, NP cells interacting with DE cells expressed *Lhx5* and *Nkx2-1*, and those interacting with NC expressed *Prtg* (Fig. 3A). These findings may underlie anterior-posterior axis inducing activities of these node derivatives. The anterior endoderm emerges very early in gastrulation to pattern the presumptive anterior neural plate, while the NC emerges later, patterning more posterior locations along the neural tube. Deconstructing positional information within the transcriptome could provide a detailed map of cells localized to axis promoting organizing centers that emerge in embryonic development. Given the role of the anterior endoderm in patterning the nascent neural plate (10) and NC for the patterning of neural tube (47–49), our identification of NP+DE and NP+NC supports the inductive profile of these cells.

A widely used approach to understand cell-communication is to use ligand-receptor pairs (16, 17). We did not find the overlap between the genes identified by a ligand-receptor analysis using CellPhoneDB (17) and the contact-specific genes identified by PIC-seq (Fig. S16). Moreover, the cell types predicted to be communicating frequently using ligand-receptor pairs do not reflect well with the anatomical structure (Fig. S17). These observations indicate that PIC-seq empowered investigation of cell contact associated gene expression which cannot be studied using ligand-receptor pairs. Furthermore, the use of co-expression of ligand-receptor pairs only shows the most frequently interacting cells and does not explain specific cells engaged in cell communication. The cell contact information by PIC-seq may provide another layer of information about cell-communication.

We developed spatial-tSNE to visualize the spatial proximity that we predicted. Previous visualization methods locate cells mainly based on transcriptomic similarities. The UMAP plot using scRNAseq data distinctly represented spatial organization of EmVE and ExVE as well as FG, MG, and HG, because their gene expression was related to their spatial interactions (Fig. 1B). However, transcriptome-based visualization could not represent the physical interaction of NP+DE, MG+NC and NP+EmVE, while these are well presented in the spatial-tSNE (Fig. 5A). Consistent with this representation, the contact-specific genes are found in association with their locations in the spatial-tSNE plot (Fig. 5B). Spatial-tSNE depends on the prediction of neighboring cell types (Figs. 3A, 3B and S12). In this work, we used 367 PICs for training. The performance of neighboring cell type prediction can be further improved as we accumulate more PICs for training. Even though spatial-tSNE cannot represent the real 3D structure of the tissue, it provides a more comprehensive map for context dependent relationships inherent in mammalian development. In addition, the computational approaches that we designed for PIC-seq can also be applied to image-based spatial transcriptomics data such as seqFISH (40) to identify contact-specific and regulated genes.

As our study is about *de novo* gene discovery, it is difficult to assign specific roles in mechanobiology, although *Gsc* is a known marker of the node, a region that has inductive properties in heterotopic grafting experiments (50).

The history of developmental biology is based on a large number of embryonic grafting experiments to define inductive interactions that occur during development (12). Grafting experiments were pivotal in our understanding of how signals produced from one cell can illicit patterning responses in another cell. However, grafting experiments are technically challenging and inherently limited to a pre-defined set of interactions. Here we take an unbiased approach to understand developmental context, producing spatial-tSNEs to provide an unbiased catalogue of potential developmental interactions. Through assessing these interactions, one could develop a comprehensive map of embryonic induction providing a set of all possible sites. While directionality can only be inferred by experiments, we present an impartial approach to study spatial gene regulation during development.

## STAR Methods

### Multiplet cell isolation

Mouse FOXA2^Venus^ embryos were collected between embryonic days (E) 7.5 and 9.5. The E9.5 embryos were dissected in order to enrich the sample with gut endoderm cells. Embryos were dissociated with Accutase (Sigma) into multiplet cells immediately after collection. The collected embryos were mixed with mouse embryonic stem cells (ESC), which were counterstained with CellVue Maroon Cell Labeling Kit (Thermofisher, # 88-0870-16) to increase the number of cells in the sample in order to avoid loss of the scarce FOXA2^Venus^ positive cells by spinning. The samples were then incubated with pre-warmed Accutase at 37^°^C. For the multiplet cell dissociation, the samples were incubated for 4 mins with 1ml of Accutase and pipetted up and down carefully to ensure cells were not dissociated into single cells. The Accutase was diluted by adding 3ml of FACS buffer 1 (10% FBS in PBS) and spun down. The cells were washed with FACS buffer 1 two more times and transferred to a FACS tube with FACS-DAPI buffer for the FACS process.

### Flow cytometry and multiplet cell capturing

Multiplet cells were isolated from FOXA2^Venus^ mouse embryos. Cells were sorted using a BD FACS Aria III. FSC/SSC gates were used to define a homogeneous population, FSC-H/FSC-W gates were used to include multiplets, remove singlets and dead cells were excluded based on DAPI inclusion. Sorting speed was kept at 100-300 events/s to eliminate sorting two or more drops containing cells into one well. Single drop deposition into the 384 well plates was verified colorimetrically based on a previously published protocol (51). Cells were sorted directly into lysis buffer containing first RT primer and RNase inhibitor, immediately frozen and later processed by MARS-seq protocol as described previously (25).

### Flow cytometry and cell sorting of NP and DE populations

Embryonic cells were isolated from E8.5 mouse embryos and incubated with cell surface antibodies specific to NP and DE cell populations. Following the incubation these populations were sorted using BD FACSAria Fusion and data was analyzed by FACSDiva 8.0.2 software as follows. FSC/SSC gates were used to define a homogeneous population, and FSC-H/FSC-A gates were used to sort singlets exclusively. For the purpose of isolating the NP cells, the embryonic cells were suspended in FACS buffer 2 (PBS, 1mM EDTA, 25mM HEPES pH 7, 2% FBS) and stained with NCAM-1:PE (Biolegend #125618), SSEA1:APC (Abcam #ab18277), and SSEA4:Alexa Fluor 488 (Thermo Fisher Scientific # 53-8843-42). NCAM-1(52)/SSEA1 (53) gate was used to select double positive populations, from which SSEA4 negative cells were selected using SSC/SSEA4 gate as described previously (53).

For the purpose of isolating the DE cells, the embryonic cells were suspended in FACS buffer 2 and stained with SSEA1:APC, SSEA4:PE (Thermo Fisher Scientific # 12-8843-42), Claudin-6:FITC (Bioss #bs-3754R-FITC), and CD24:BV421 (BD Horizon™ #562563). Here, CD24(54)/Claudin-6 (55) gate was used to select double positive cells followed by the selection of populations negative for SSEA1 and SSEA4 using the SSEA1/SSEA4 gate to avoid contamination with NP and visceral endoderm cells (53, 54). Isolated NP and DE cells were maintained in culture using the StemFlex Medium (Thermo Fisher) at 37°C, 5% CO2 incubator.

A sub population of the DE cells were infected with 10^9^ TU/ml of GFP comet-pD2109-CMV lentiviral particles (ATUM) and 5 μg/ml of polybrene. After 24h infection, GFP positive DE cells were selected by GFP expression using BD FACSAria. Following a 48 hr co-culture of GFP+ DE and GFP-NP cells in a 1:1 ratio, the BD FACSAria was used to sort the GFP positive DE cells and GFP negative NP cells according to their GFP expression using SSC/GFP gate.

### DE and NP contact-specific genes quantification by RT-qPCR

We isolated total RNA from DE and NP cells individual cultures and co-cultures using the PureLink RNA Mini kit, as per manufacturer instruction and eluted total RNA in 50 μL RNase/DNase-free H_2_O. Then, we reverse-transcribed to cDNA 10 ng of total RNA using Superscript Vilo cDNA synthesis kit. Finally, we performed real-time PCR (qPCR) in QuantStudioTM 5 (Applied Biosystems) using PowerUp SYBR Green master mix (Thermo Fisher Scientific) and the following reaction conditions. The initial denaturation step was performed at 95 °C for 2 min, followed by 40 cycles of 95 °C for 15 s and 60 °C for 60 s. We used the comparative CT method (ΔΔCt) to quantify relative gene expression, normalizing the expression of our target genes with the housekeeping gene *Gapdh*. All samples were run using the following primers: *Gapdh*: 5′-CATCACTGCCACCCAGAAGACTG-3′ (F) and 5′-ATGCCAGTGAGCTTCCCGTTCAG-3′ (R); *Lhx5*: 5’-CTCGACCGCTTTCTGCTGAA-3’ (F) and 5′-CGCTCGGAGAGATACCTTGC-3′ (R); *Nkx2-1*: 5’-AGGACACCATGCGGAACAG-3’ (F) and 5′-CCATGCCGCTCATATTCATGC-3′ (R); *Gsc*: 5’-GACGAAGTACCCAGACGTGG-3’ (F) and 5′-CGGTTCTTAAACCAGACCTCCA-3′ (R); *Rax*: 5’-TGGGCTTTACCAAGGAAGACG-3’ (F) and 5′-GGTAGCAGGGCCTAGTAGCTT-3′ (R)

### Data processing of scRNAseq and PIC-seq data of mouse embryo

All scRNAseq libraries were sequenced using Illumina NextSeq 500. Sequences were mapped to mouse mm9 genome, de-multiplexed and filtered as previously described (25, 26). We estimated the level of spurious unique molecular identifier (UMIs) in the data using statistics on empty MARS-seq wells as previously described (25). Mapping of reads was done using HISAT (version 0.1.6) (56); reads with multiple mapping positions were excluded.

Among the transcriptome of 6,600 single cells and 382 PICs, we filtered the low-quality samples that had UMI counts over 2^17^ or less than 256 2^8^. The remaining samples contained 6,288 single cells and 367 PICs. To remove any batch effect, we used the Seurat v3 standard integration workflow (57). The 13 clusters were obtained using a graph-based Louvain clustering algorithm.

### SVMs for classification

For the classification of PICs, we trained a multiclass classifier for SVMs using a MATLAB (version R2020a) function ‘fitcecoc’. The classifier consists of multiple SVM binary learners in a one-vs-one design. We trained three SVMs for each stage (E7.5, E8.5, and E9.5) of PIC-seq and scRNAseq data. We used the major clusters for each stage (DE, N, ExVE, EmVE, PS, PE for E7.5; NP, FG, DE, MG, HG, NC for E8.5; FG, MG, HG, L, NC for E9.5) and the top 5 DEGs for each cluster. We run 10-fold cross-validation for the trained models using a MATLAB function ‘crossval’ and calculated the error rate using ‘kFoldLoss’ with 10. We calculated significance of the frequencies of PICs against doublets identified from scRNAseq using Fisher’s exact test.

### SVMs for neighboring cell type prediction

For the prediction of the identity of neighboring cells, we applied multiclass SVM using ‘fitcecoc’ function in MATLAB R2020a. We only used the contact-specific genes for training and prediction. The SVMs for each cell type were trained using the PIC-seq data with their annotations. For example, to train a SVM for NP, PICs included in NP+NP, NP+DE, and NP+NC were used. To predict the neighboring cell type from the single cell transcriptome, we made artificial PIC-seq by mixing the transcriptome of the cell of interest with all other cells one by one. The artificial PIC-seq data for the cell of interest were predicted using the corresponding SVM. The voting schemed (i.e., most frequent) is used to assign the neighboring cell type. We applied this scheme for all single cells.

### Statistical analysis and enrichment analysis

The p-value for cell-type specific and contact-specific marker genes was calculated by using a two-sided Wilcoxon rank sum test. The p-value for RT-qPCR of contact specific genes was calculated by using a two-sided Student’s t-test. We used Enrichr (39), an enrichment analysis tool, to investigate the enriched GO terms and KEGG pathways for each marker gene group.

### Staining of mouse embryo

E8.5 embryos were isolated in PBS and then fix overnight in 4% paraformaldehyde (PFA) overnight at 4 °C. The following day, embryos were washed in PBT (PBS containing 0.1% Tween-20), dehydrated in an ascending methanol sequence, xylene treated, embedded in paraffin, and sectioned at 6.5μm. Immunofluorescence was performed on 6.5μm deparaffinized sections. Sections were subjected to antigen retrieval in Tris buffer pH 10.2 for 10 min, washed in 0.1% PBT and incubated in blocking buffer (0.5% milk powder, 99.5% PBT) for 2 h at room temperature. Primary antibodies were incubated in blocking buffer overnight at 4 °C. The following day, the sections were washed three times with PBT and incubated for 1 h with corresponding secondary antibodies in blocking buffer at room temperature. After three washes in PBT, DAPI (Sigma-Aldrich, 1:2000) was added to counter-stain the nuclei. The sections were mounted using Prolong Gold Antifade Reagent (Invitrogen, P36934) and imaged using Zeiss LSM-700 confocal microscope. The following primary antibodies were used: FOXA2 (Thermofisher, 7H4B7, 1:250), SOX2 (eBioscience, 14-9811-80, 1:250), NKX2-1 (Abcam, ab76013, 1:250). Secondary antibodies were Alexa Fluor conjugates 488, 555, and 647 (Life Technologies) at 1:500.

### spatial-tSNE

Spatial-tSNE is designed for visualization of neighboring cells in scRNAseq data, which is modified from Tsne (58) by considering neighboring cell information. In the original tSNE, the 2D embeddings is obtained by rearranging each cell based on the probability of the pairwise similarities of the cells on the original high-dimensional space. The pairwise similarity of two cells *i* and *j* in the high-dimensional space is defined by the probability, 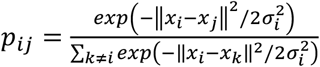 where *x*_*i*_ is the data points of cell *i* and σ_*i*_ is the variance of the Gaussian which is centered on data point *x*_*i*_. The distance between two cells in the reduced t-SNE dimension is determined by the probability *p*_*ij*_ in high-dimensional space.

Spatial-tSNE shows the clustering and spatial information at the same time by changing the similarity probability for the cells that are predicted to neighbor with each other. To reflect spatial information in the t-SNE dimension, we modified the probability *p*_*ij*_ so that the similarity probability of two neighboring cells is the maximum probability of all pairs. With these modified probabilities, the predicted neighbor cells are located at the border of the two clusters without ruining the relative position of other cells. To clearly visualize neighbored cells, an imaginary line was drawn between the two populations, which can be rotated so that cells have the longest distance to it. Other cells are re-arranged based on their probabilities.

## Data and code availability

Source code and input files for the PIC classification and the neighboring cell type prediction are available at https://github.com/neocaleb/NicheSVM. Source code for spatial-tSNE is available at https://github.com/wgzgithub/sp_tSNE. The PIC-seq data can be downloaded from Gene Expression Omnibus database with accession number (GSE182393).

## Acknowledgments

The Novo Nordisk Foundation Center for Stem Cell Medicine is supported by a Novo Nordisk Foundation grant (NNF21CC0073729 and NNF17CC0027852). This work is also supported by Lundbeck Foundation (R324-2019-1649, R313–2019–421) to KJW, by Swiss Cancer League, Seeds of Science, UAMS, SNSF, Fundação para a Ciência e a Tecnologia, Fundamental Mandates (Stichting tegen Kanker—Fondation contre le Cancer) to R.G. The National Research Foundation of Korea (NRF) grant funded by the Korea government (MSIT) to JIK (No. 2021R1F1A1063914 and 2021M3H9A2096988), La Caixa Funding (LCF/BQ/PR20/11770006) to EM and Lundbeck Foundation (R198-2015-412), Fundação para a Ciência e a Tecnologia Grant 2020.05319.BD to A.M.P, Thelma Newmeyer Corman Chair in Cancer Research, the VCU Commercialization Fund, NIH grants NIH/NCI R01CA259599 and R01CA244993 to P.B.F., and Independent Research Fund Denmark (8020-00100B and 6110-00009) to JMB.

