## Supplementary Materials for "Neighbor-specific gene expression revealed from physically interacting cells during mouse embryonic development"

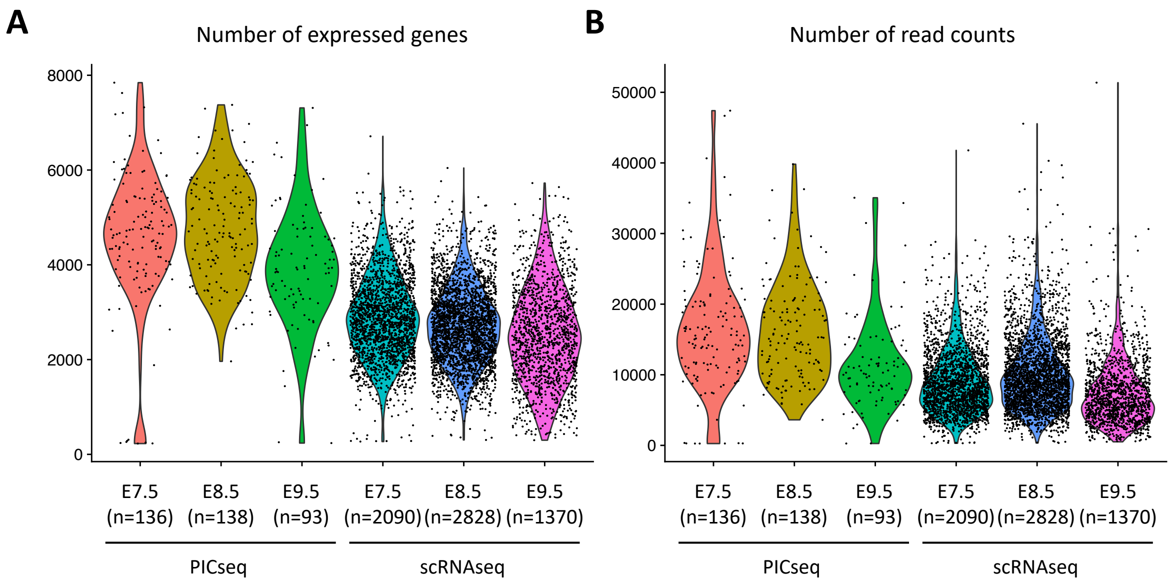


**Figure 1. The quality control of PIC-seq and scRNAseq data of the mouse embryo at E7.5-E9.5.** (**A**) The number of expressed genes and (**B**) the number of read counts in each PIC and cell for each sample.

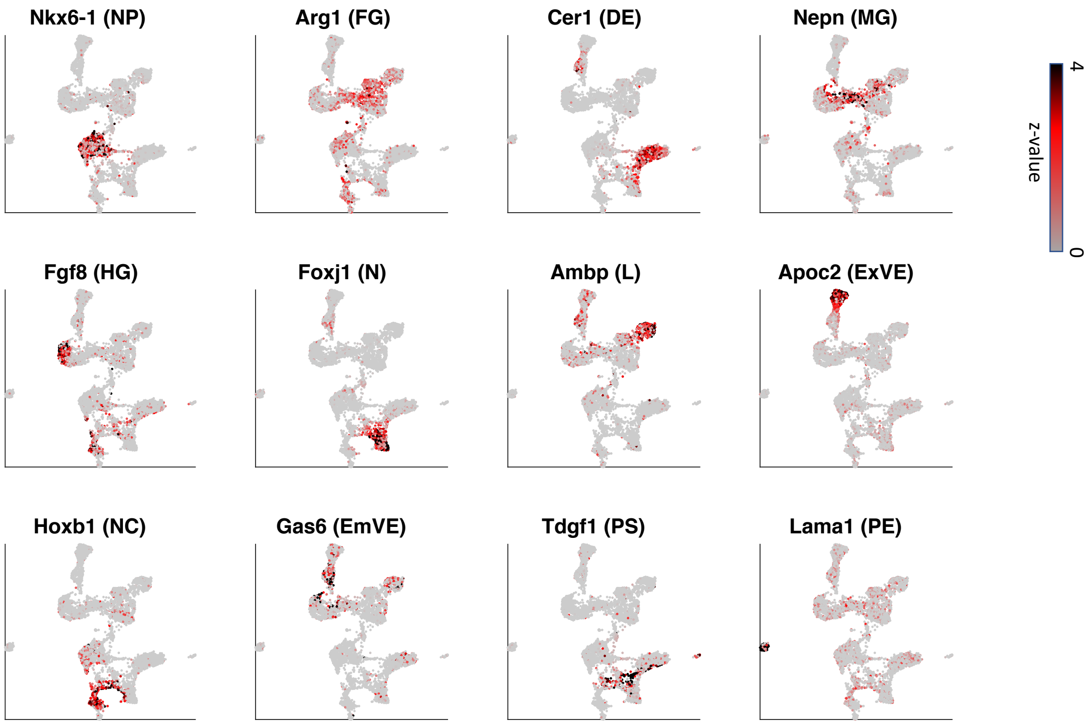


**Figure S2.** Expression profiles of 12 cell-type marker genes on a two-dimensional plot. NP: Neural progenitors, FG: Foregut, DE: Definitive endoderm, MG: Midgut, HG: Hindgut, N: Node, L: Liver, ExVE: Extra-embryonic visceral endoderm NC: Notochord, EmVE: Embryonic visceral endoderm, PS: Primitive streak, PE: Parietal endoderm. The expression levels of a marker gene are shown for the corresponding cell type.

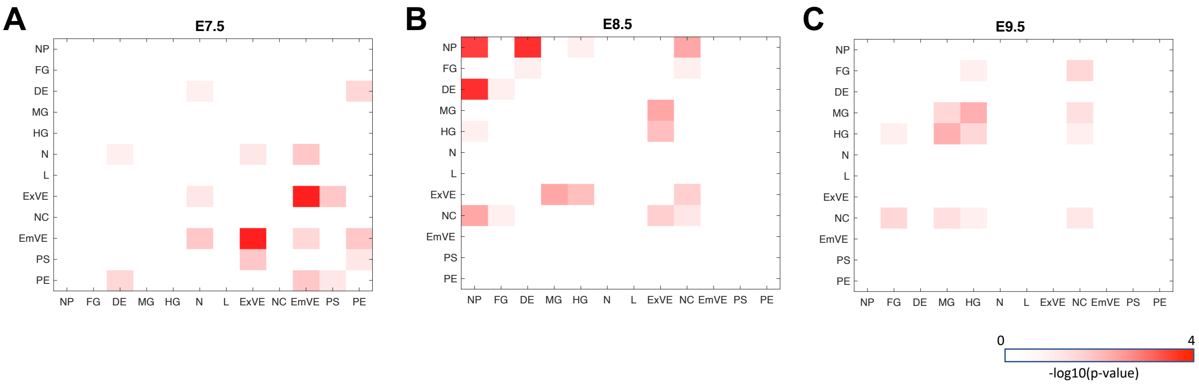


**Figure S3.** The significance of the frequencies of co-occurring cell types in PIC-seq compared with erroneous doublets found in scRNAseq at (**A**) E7.5, (**B**) E8.5 and (**C**) E9.5. The identified interactions from PIC (e.g. ExVE+EmVE at E7.5 and NP+DE at E8.5) were significantly more frequent than the erroneous doublets.

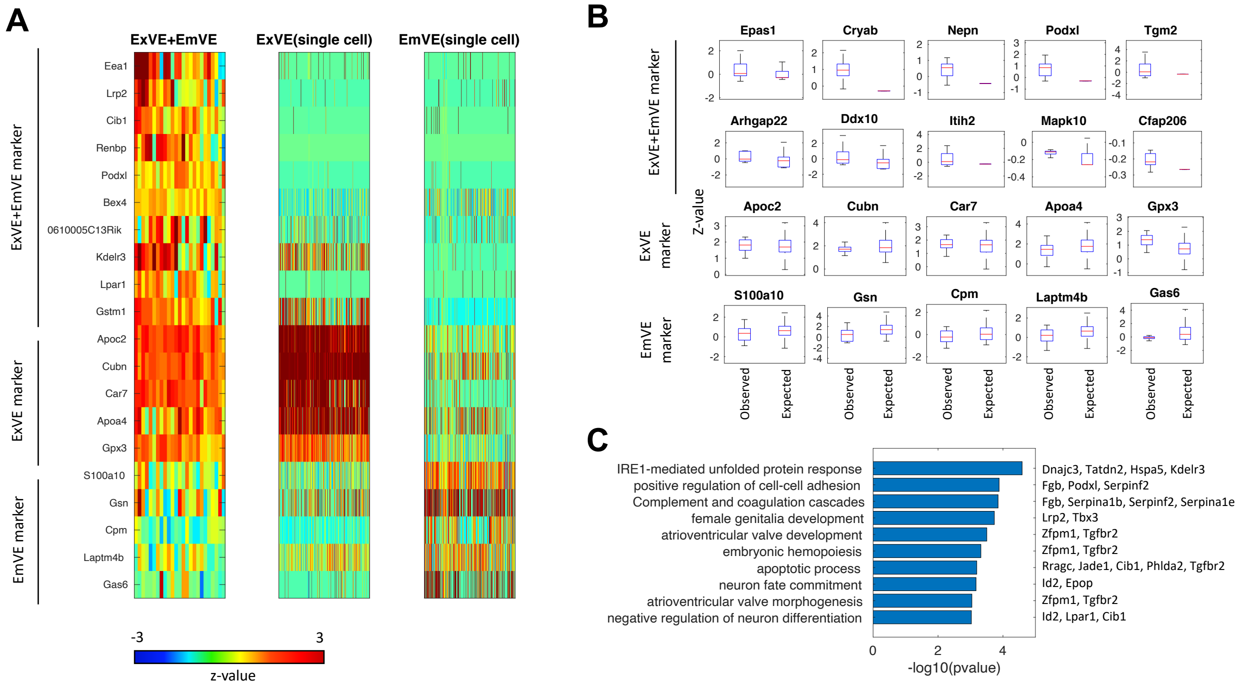


**Figure S4. PIC-seq analysis identified genes highly expressed in ExVE+EmVE.** (**A**) A heatmap of PIC-seq for ExVE+EmVE showed contact specific expression as well as the marker genes for each cell type. (**B**) The expression levels for the contact specific genes are significantly higher in the PIC-seq compared with expected expression levels obtained from scRNAseq. (**C**) GO and KEGG pathway terms enriched in the contact-specific genes for ExVE+EmVE.

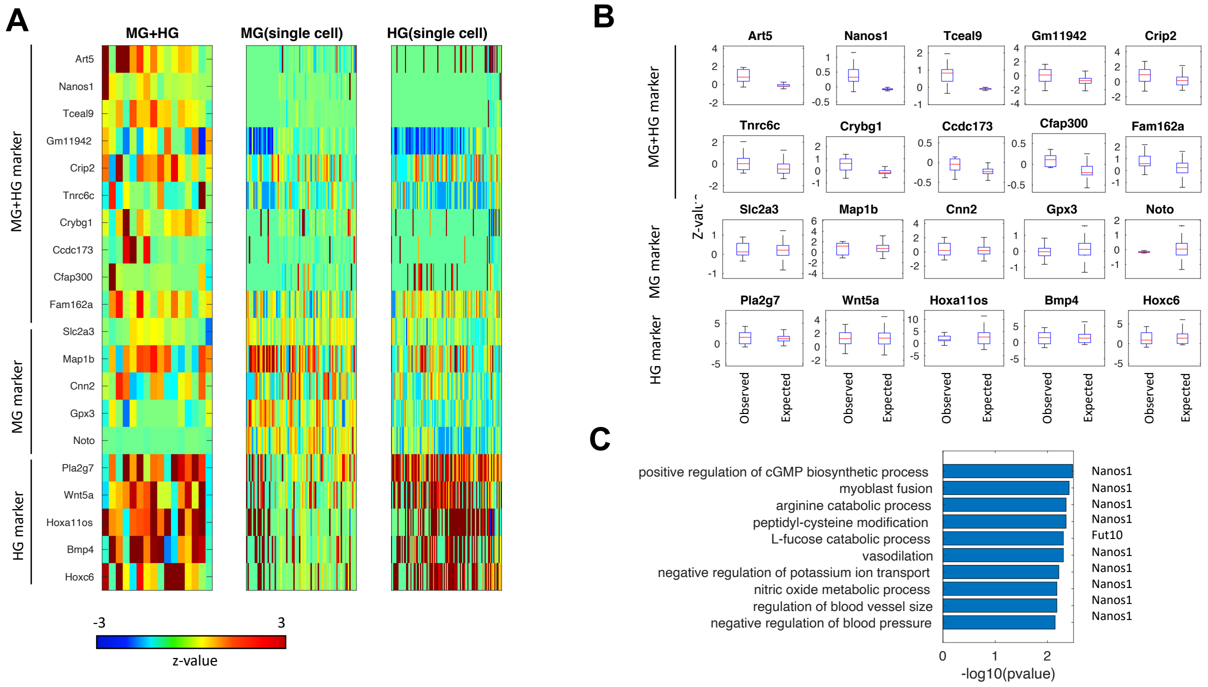


**Figure S5. PIC-seq analysis identified genes highly expressed in MG+HG.** (**A**) A heatmap of PIC-seq for MG+HG showed contact specific expression as well as the marker genes for each cell type. (**B**) The expression levels for the contact specific genes are significantly higher in the PIC-seq compared with expected expression levels obtained from scRNAseq. (**C**) GO and KEGG pathway terms enriched in the contact-specific genes for MG+HG.

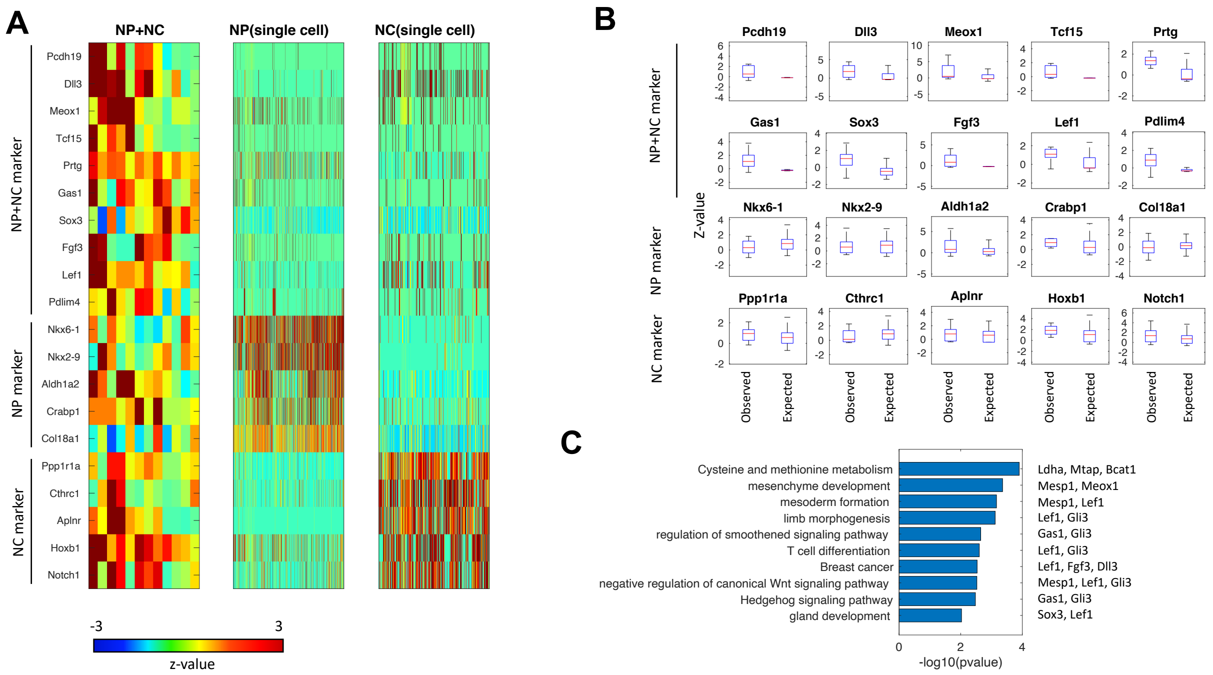


**Figure S6. PIC-seq analysis identified genes highly expressed in NP+NC.** (**A**) A heatmap of PIC-seq for NP+NC showed contact specific expression as well as the marker genes for each cell type. (**B**) The expression levels for the contact specific genes are significantly higher in the PIC-seq compared with expected expression levels obtained from scRNAseq. (**C**) GO and KEGG pathway terms enriched in the contact-specific genes for NP+NC.

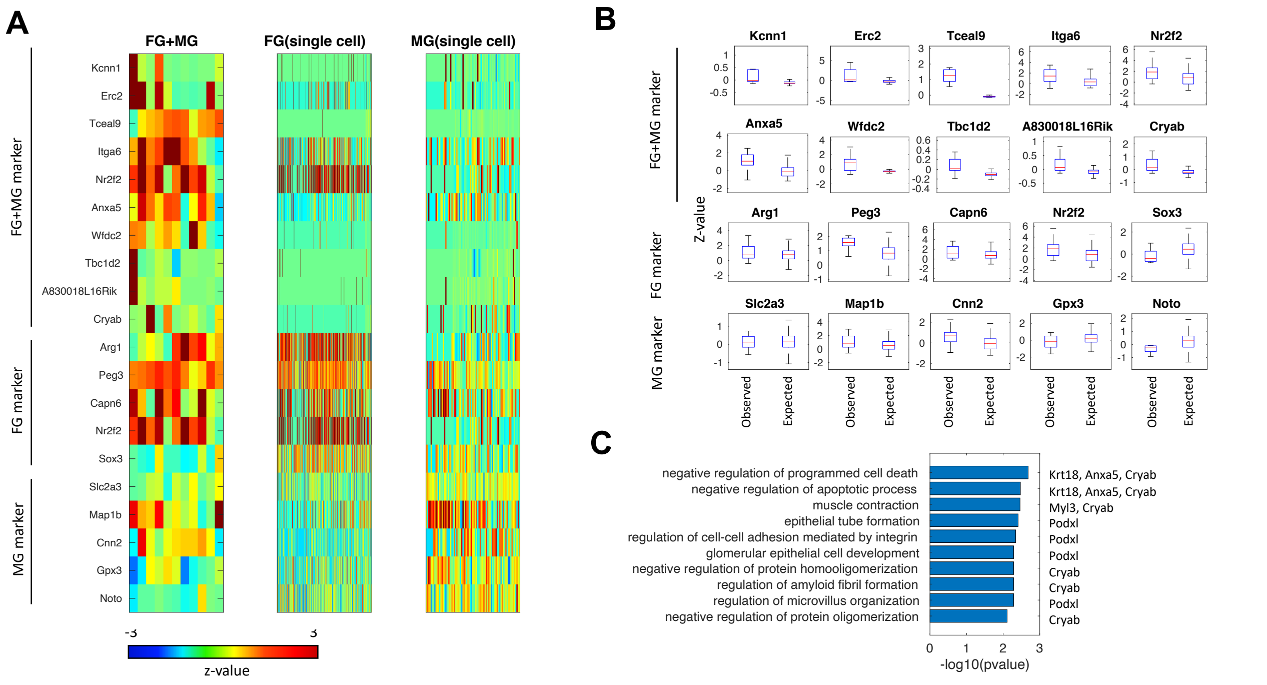


**Figure S7. PIC-seq analysis identified genes highly expressed in FG+MG.** (**A**) A heatmap of PIC-seq for FG+MG showed contact specific expression as well as the marker genes for each cell type. (**B**) The expression levels for the contact specific genes are significantly higher in the PIC-seq compared with expected expression levels obtained from scRNAseq. (**C**) GO and KEGG pathway terms enriched in the contact-specific genes for FG+MG.

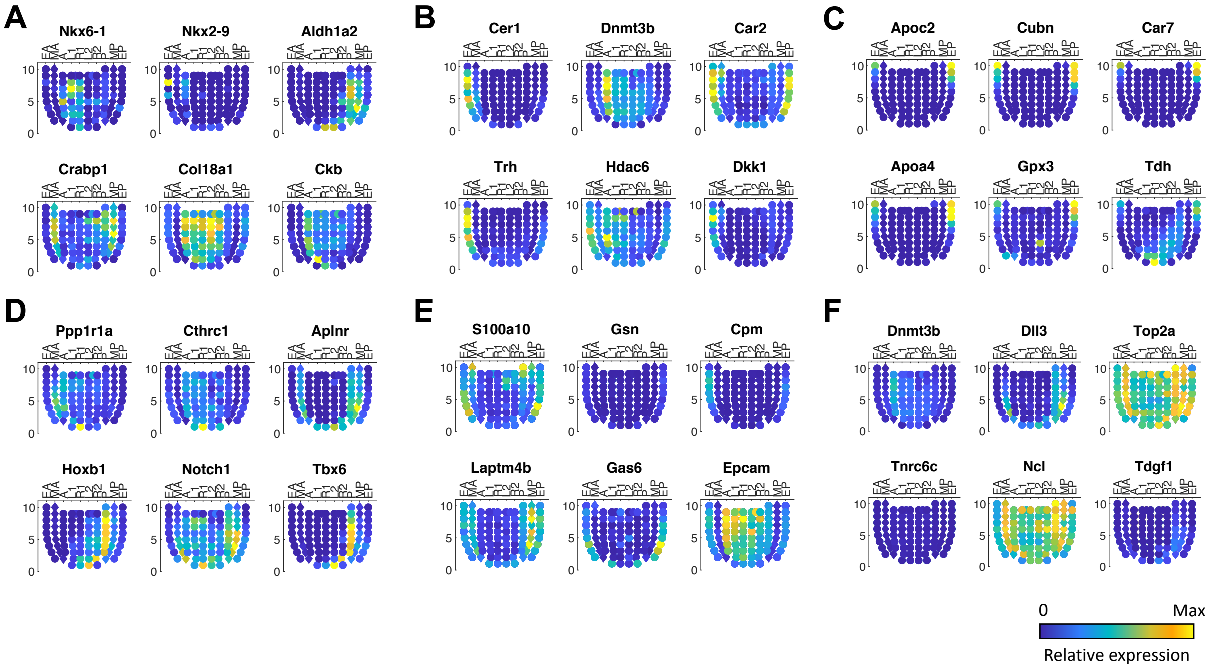


**Figure S8. Expression levels of top 6 marker genes on Geo-seq data for each cell type** (**A**) NP, (**B**) DE, (**C**) ExVE, (**D**) NC, (**E**) EmVE, (**F**) PS. While some of these markers are not expressed in the correct tissue, we assume that this is because the marker sets span E7.5-9.5, whereas these corn plots are based on data from E7.5 embryos. EA: Anterior Endoderm; MA: Anterior Mesoderm; A: Anterior; L1: Anterior Left Lateral; R1: Anterior Right Lateral; L2: Posterior Left Lateral; R2: Posterior Right Lateral; P: Posterior; MP: Posterior Mesoderm, EP: Posterior Endoderm.

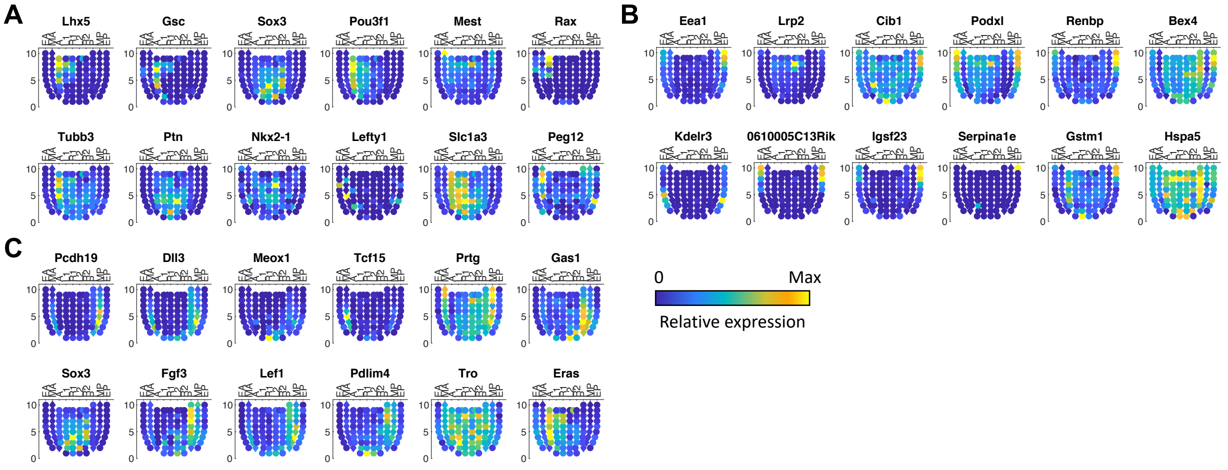


**Figure S9. Expression levels of top 12 contact specific marker genes on Geo-seq data**. (**A**) NP+DE, (**B**) ExVE+EmVE, and (**C**) NP+NC marker genes.

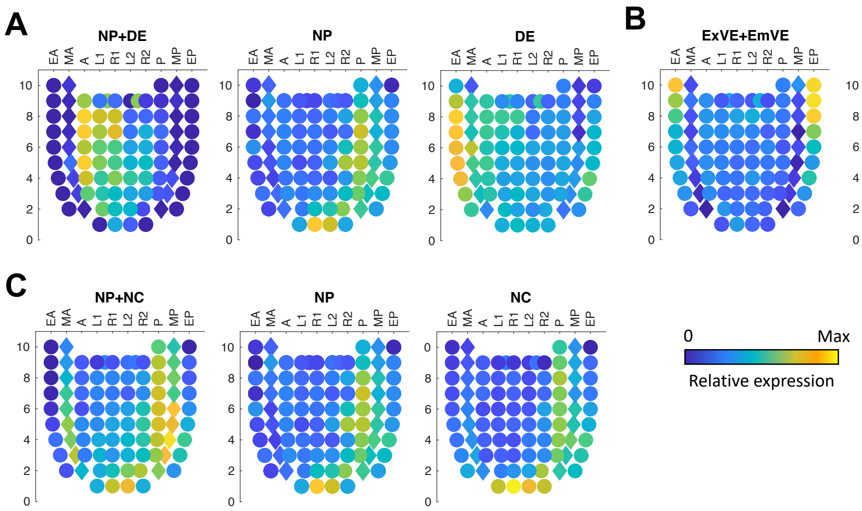


**Figure S10. Average expression levels of contact specific marker genes (Supplementary Table S2) on Geo-seq data**. (**A**) NP+DE, (**B**) ExVE+EmVE, and (**C**) NP+NC.

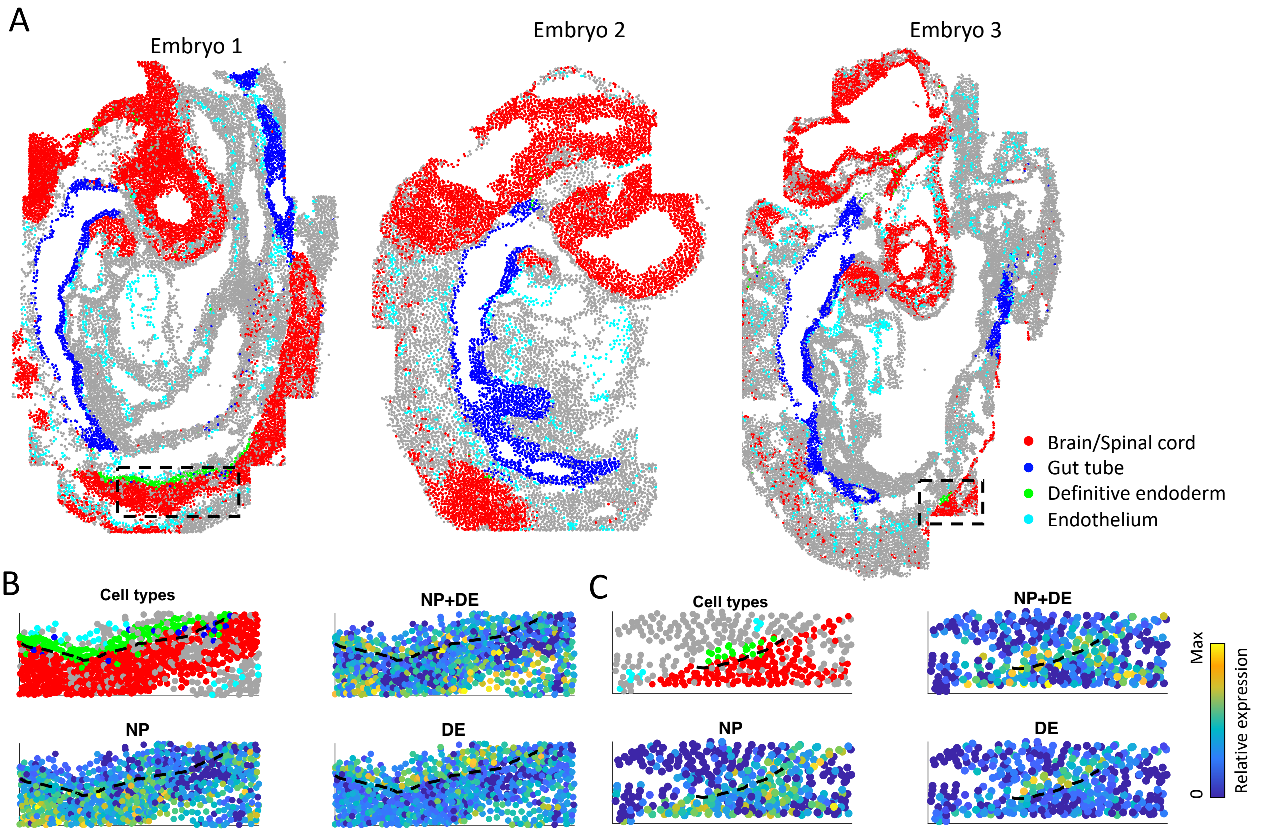


**Figure S11. Anatomical position of cell types and average expression profile of contact-specific genes in mouse embryo E8.5-E8.75 on seqFISH data**. (**A**) Anatomical position of cell types (Brain/Spinal cord (or NP), Gut tube, Definitive endoderm, and Endothelium) in the mouse embryo (**B-C**) Average expression profile of contact-specific and cell type-specific genes (NP+DE, NP, and DE). The boxed region in A is shown at higher magnification in each panel of B and C, with the dotted line showing the interface between the two cell types and the expression of the contact specific genes is apparent along this border

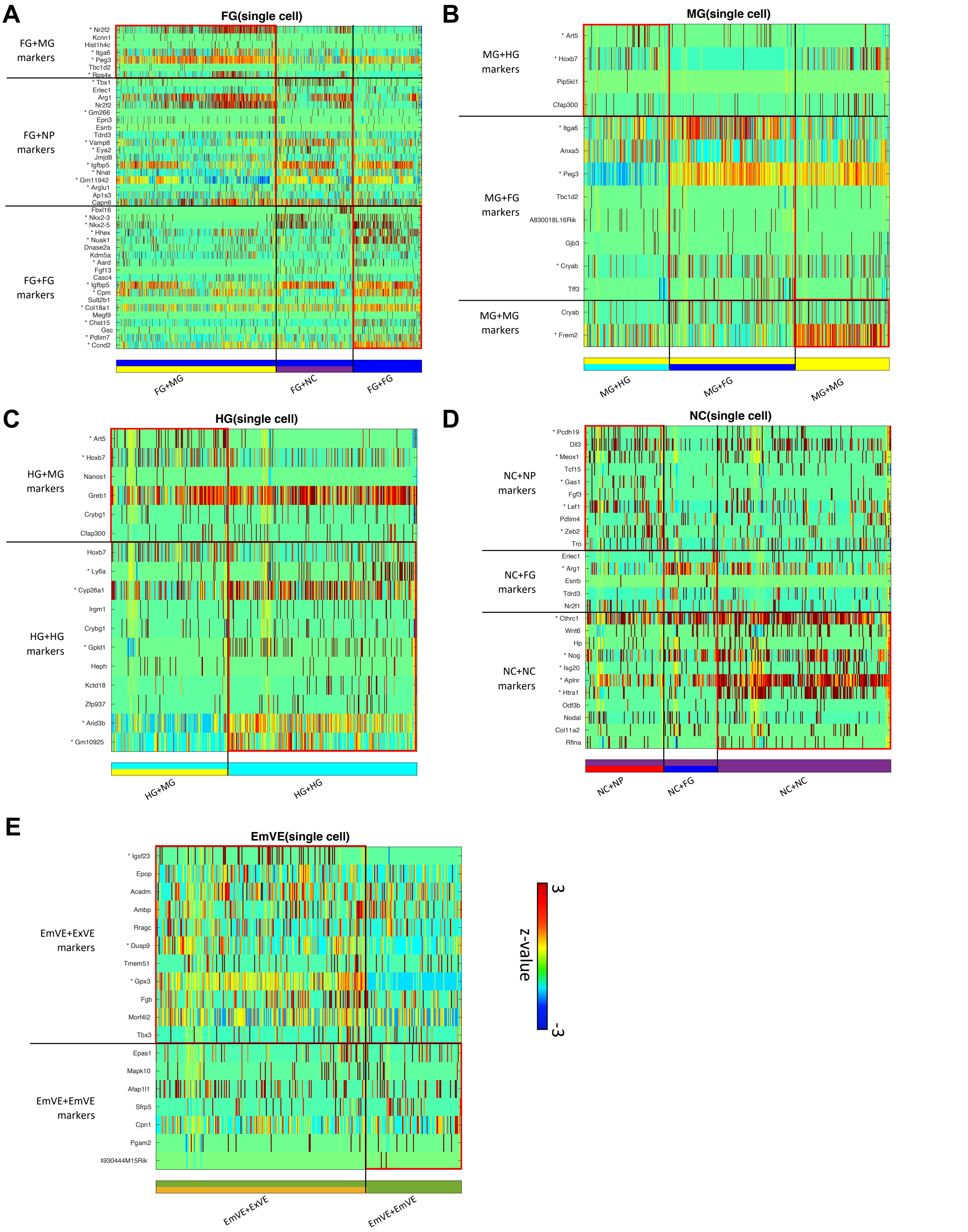


**Figure S12. Prediction of the neighboring cell type and the contact-specific expression.** (**A**) FG cells predicted to neighbor with MG, NC, and FG, (**B**) MG predicted to neighbor with HG, FG, and MG, (**C**) HG predicted to neighbor with MG, and HG, (**D**) NC predicted to neighbor with NP, FG, and NC, (**E**) EmVE predicted to neighbor with ExVE and EmVE. *: Wilcoxon rank sum test p-value<0.01

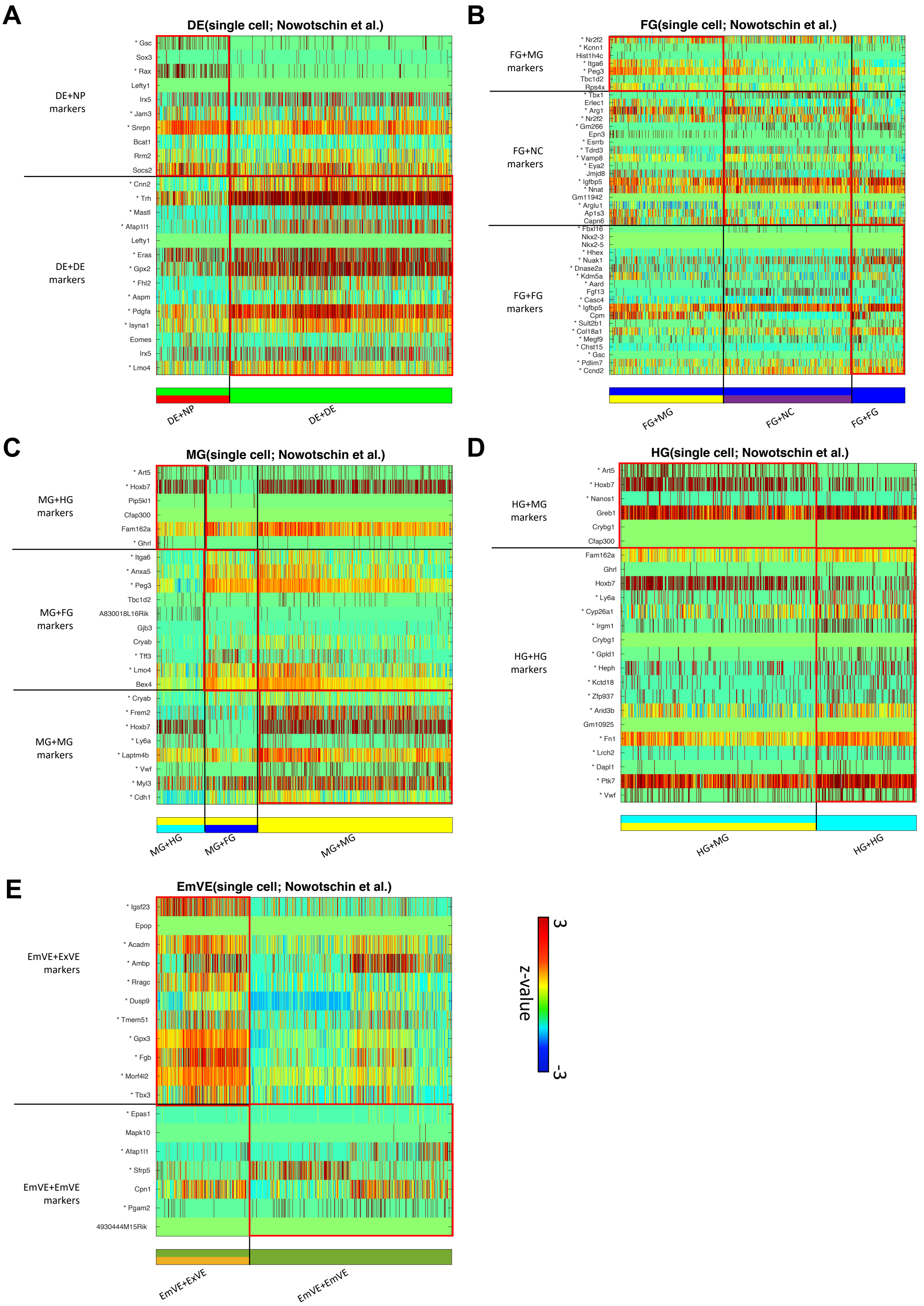


**Figure S13. Prediction of the neighboring cell type using an independent publicly available scRNAseq dataset of mouse endoderm during E3.5-E8.75 (Nowotschin *et al.*).** (**A**) DE, (**B**) FG, (**C**) MG, (**D**) HG, (**E**) EmVE. *: Wilcoxon rank sum test p-value<0.01.

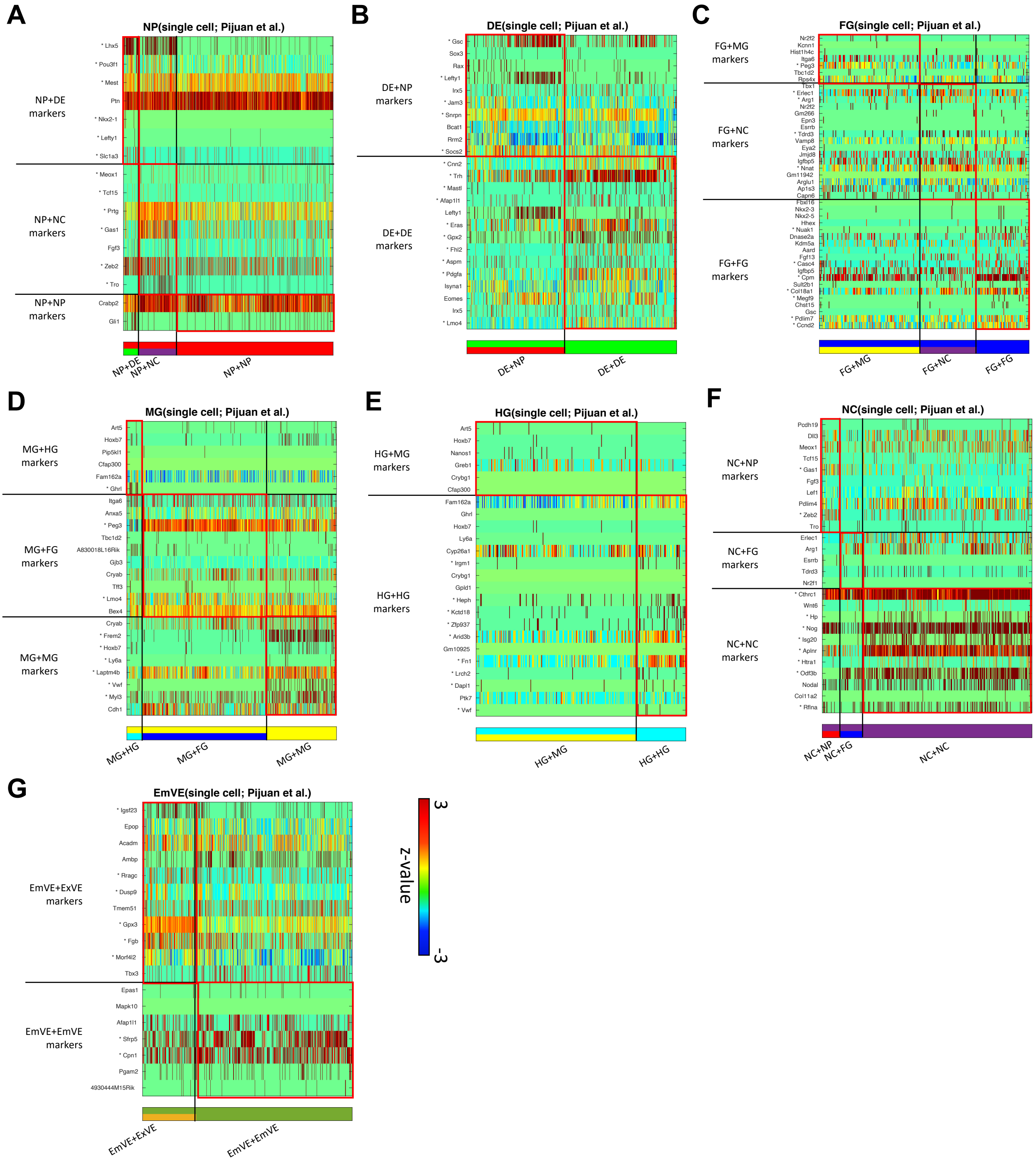


**Figure S14. Prediction of the neighboring cell type using an independent publicly available scRNAseq dataset of mouse endoderm during E3.5-E8.75 (Pijuan *et al.*).** (**A**) NP, (**B**) DE, (**C**) FG, (**D**) MG, (**E**) HG, (**F**) NC, (**G**) EmVE. *: Wilcoxon rank sum test p-value<0.01.

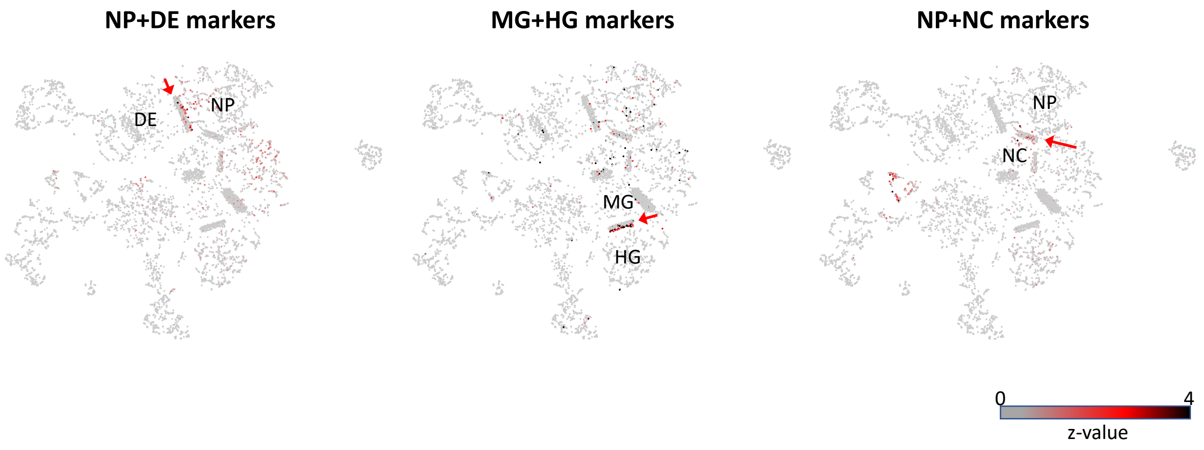


**Figure S15. Contact specific marker gene expression on spatial-tSNE.** A spatial tSNE plot recapitulates the spatial distribution of cells. The arrows indicate the cells predicted to locate in the border of the corresponding two cell types.

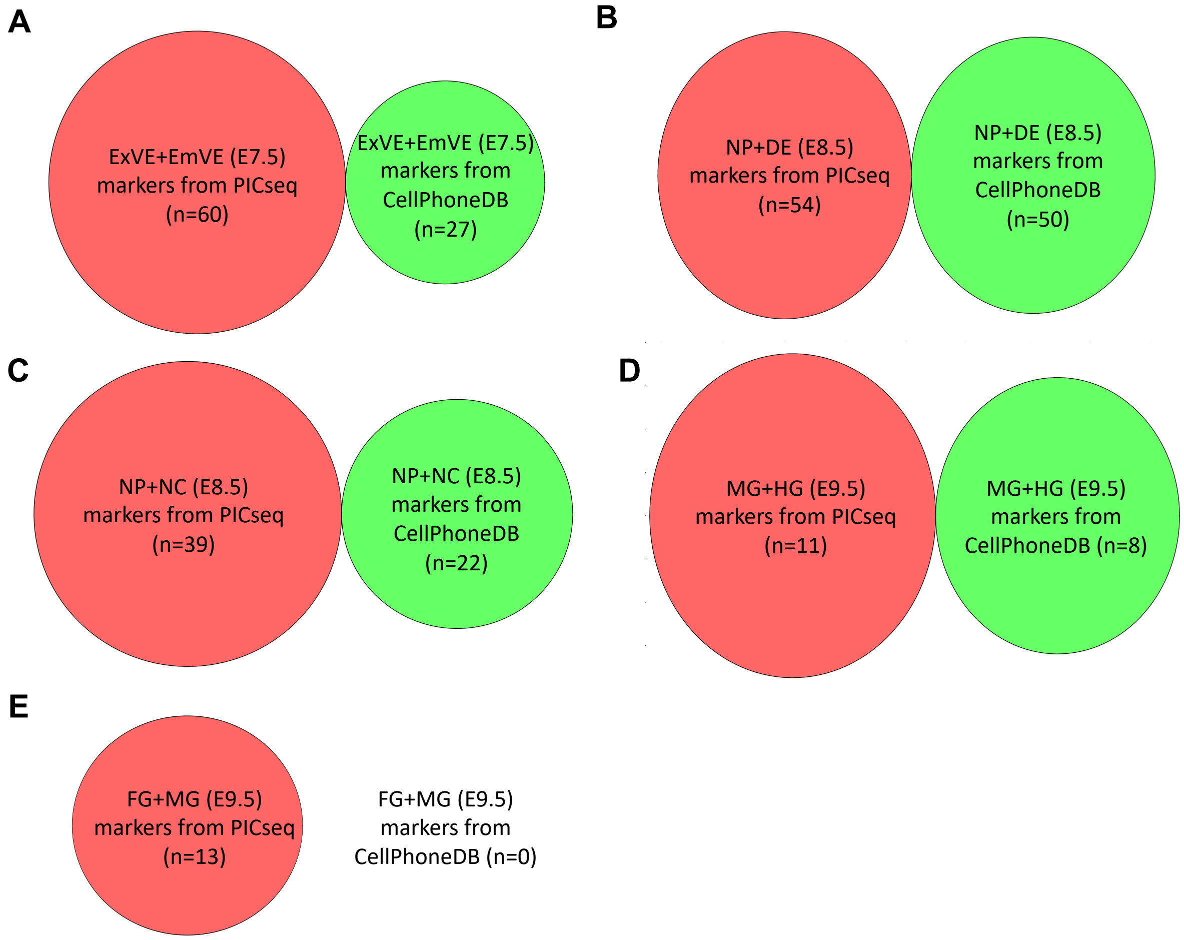


**Figure S16. Discrepancy between contact-specific genes identified by PIC-seq and gene sets identified a ligand-receptor analysis CellphoneDB.** (**A**) ExVE+EmVE, (**B**) NP+DE, (**C**) NP+NC, (**D**) MG+HG, (**E**) FG+MG.

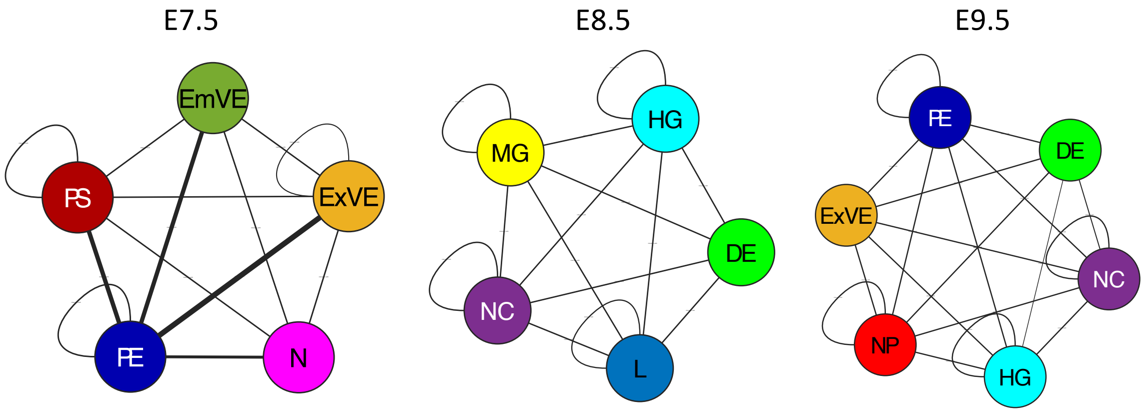


**Figure S17. The frequently communicating cell type predicted by CellphoneDB against scRNAseq at E7.5, E8.5 and E9.5.** The width of the line represents relative frequencies. The prediction using the ligand-receptor pairs cannot tell the specific cell types interacting with each other.
